# Modeling lung cell development using human pluripotent stem cells

**DOI:** 10.1101/2021.07.16.452691

**Authors:** Shuk Yee Ngan, Henry Quach, Joshua Dierolf, Onofrio Laselva, Jin-A Lee, Elena Huang, Maria Mangos, Sunny Xia, Christine E. Bear, Amy P. Wong

## Abstract

Human pluripotent stem cells (hPSC) differentiations can capture developmental phenotypes and processes. They are useful for studying fundamental biological mechanisms driving tissue morphogenesis and cell lineage development. Here, we show temporal development of lung cell lineages using hPSC that recapitulate developmental milestones observed in primary tissue, the generation of renewable fetal lung epithelial spheroids, and the functional utility of the lung models at different differentiation stages for cystic fibrosis disease modeling. We first show the presence of hPSC-derived lung progenitor cells reminiscent of early trimester lung development and containing basal stem cells that generate renewable airway spheroids. Maturation and polarization in air liquid interface (ALI) generates additional epithelial cell lineages found in adult airways, including pulmonary neuroendocrine, brush, mature basal, ciliated and secretory cell types. Finally, pseudotime and RNA velocity analyses of the integrated datasets from fetal and ALI stages reveal both previously identified and new cell lineage relationships. Overall, hPSC differentiation can capture aspects of human lung development and potentially provide important insight into congenital causes of diseases.

## INTRODUCTION

Human lung organogenesis begins around four weeks of gestation (GW) as a small batch of cells emerging from the foregut endoderm. Co-development of the lung epithelium and the surrounding mesenchyme is facilitated by mutual cross-talk between these two cell types, which ultimately guides the formation of an intricate airway tree^1, 2^. During the pseudoglandular stage (GW5-17), formation of the major airways, respiratory parenchyma and other non-epithelial cell types contribute to the developing organ. Proximal-distal patterning during this stage leads to the first signs of epithelial lineage differentiation, forming basal, ciliated, neuroendocrine (NE) and secretory cells. As the lung develops in the canalicular stage (GW16-27), additional cellular differentiation occurs with the formation of the distal lung and emergence of primitive alveolar cells. The saccular stage (GW24-38) marks the expansion of air sacs and the production of surfactants in preparation for the lungs to switch to an air breathing organ after birth^3^. The final alveolar stage (GW24 up to 10 years) begins shortly before birth and continues well into childhood years with maturation of the alveoli and expansion of the capillaries, nerve and gas exchange regions of the lungs.

Single cell RNA sequencing (scRNA-seq) has enabled our recent understanding of the complexities of the cellular phenotypes that make up the human and mouse lungs^4–9^. Importantly, the identification of new cell types or cell states have been implicated in human lung diseases^5, 7, 10^ and the question of the origins of these cell types have sparked renewed interest in the understanding of human lung development^11, 12^. New *in-vitro* model systems from isolated human fetal lung or hPSC-derived tissues are shedding light on key pathways regulating cell lineage formation through experimental manipulation^11, 13–15^. Pluripotent stem cells are derived from the inner cell mass of the pre-implantation embryo (called embryonic stem cells, ESCs), or are artificially generated through reprogramming of somatic cells (called induced pluripotent stem cells, iPSCs)^16, 17^. Early step-wise iPSC differentiation protocols modeling developmental milestones based largely on mouse lung development showed that bona fide lung epithelial tissues can be generated^18–20^ and used to model cystic fibrosis airway disease^18, 21–23^. Modifications in differentiation protocols and identification of cell surface molecules for fluorescence activated cell sorting (FACS) of putative progenitor cells have significantly enriched for cells capable of forming 3-dimensional lung epithelial spheroids for modeling disease^13, 14, 24, 25^ and fetal development^11, 26^.

Human PSC-derived lung models are also currently paving the way for the identification of new drugs to treat individual cystic fibrosis (CF) disease^13, 14, 18, 22^. Indeed iPSC derived from CF individuals have been shown to be a valuable asset for assessing new emerging cystic fibrosis transmembrane conductance regulator (CFTR) modulators (small molecules aimed at targeting the defects caused by genetic mutations) at a personalized level^23, 27, 28^. We previously showed a directed differentiation protocol to generate robust CFTR expressing cells from human PSC and importantly, how CF iPSC-derived airway epithelia could be used to model CF disease *in vitro*^18, 22^. Since then, others have demonstrated the use of CF iPSC-derived 3D airway spheroids in forskolin-induced swelling (FIS) assays in the presence of CFTR modulator compounds as a method for testing the efficacy of the compounds at a patient-specific level^14^, similar to primary rectal organoids^29^. However, the long process of generating sufficient lung tissues from iPSC for rapid and high content screens is cumbersome and costly. The ability to generate another source of cells that express functional CFTR for rapid testing of CFTR modulator effectiveness can be beneficial.

Here we show an improved hPSC differentiation method to generate airway cells that model human lung cell development. We employed scRNA-seq to determine the cellular heterogeneity and the developmental course of the cells *in vitro*. We validated the phenotypic and functional characteristics of the hPSC-derived lung cells throughout the differentiation stages and correlated the expression patterns of hPSC-derived fetal lung cells to human pseudoglandular fetal lung tissues. We also show that fetal lung epithelial cells (EC) isolated by magnetic-activated cell sorting (MACS) for the epithelial cell adhesion molecule, CD326^+^, form 3D expandable epithelial spheroids containing multipotent basal stem cells. These organoids can undergo multilineage differentiation in defined media or air liquid interface (ALI) culture. Finally, we show a developmental connection between cells that emerge during the differentiation process and highlight unique cell lineage relationships from the fetal to mature cell states. This may provide important clues for future cell engineering platforms in tissue regeneration and identify molecular regulators of airway cell differentiation during homeostasis, development and disease.

## RESULTS

### Stage-specific characterization of *in-vitro* hPSC-derived fetal lung cells

We previously demonstrated a directed differentiation method to generate proximal airway epithelia expressing functional CFTR for CF disease modeling^18, 22^. Integrating new developments in directed differentiation protocols toward lung cell lineages^11, 15^, we have refined our previous protocol to generate robust airway epithelial cell lineages (**Fig. 1**) mimicking a developmental trajectory of the cells as they differentiate and mature *in-vitro*. To demonstrate the efficiency of the new protocol, we directly compare both old (*Wong 2015*) and new protocols in subsequent analyses using two human pluripotent stem cell lines (human embryonic stem cell line CA1^18, 22^ and an iPSC lung reporter line NKX2-1^GFP14, 15^) previously used in development of differentiation protocols. Using a commercially defined kit to generate highly efficient definitive endoderm (>95%), cells are pushed towards anterior ventral foregut endoderm differentiation following a brief dual SMAD inhibition (transforming growth factor beta (TGFβ) and bone morphogenetic protein 4 (BMP4) inhibition) period (2 days)^30^ in media containing supplements favoring anterior foregut ventral endoderm (AFVE) differentiation. At the end of this stage, >90% of the cells are sex determining region Y-box-2 (SOX2) positive with no detectable midgut pancreatic and duodenal homeobox-1 (PDX1^+^) and low level of hindgut caudal type homeobox-2 (CDX2^+^) cells (4.2%) (**Supplementary Fig. 1A**). Gene expression analysis of foregut endoderm shows marked up-regulation of foregut markers forkhead box G1 (*FOXG1)*, *SOX2,* and modest elevation in the earliest lung marker NK2 Homeobox 1 (*NKX2-1*) (**Supplementary Fig. 1B**). Compared to fetal control RNA (whole fetal lung from GW18 tissue), airway epithelial specific lineage markers were not highly expressed by bulk gene expression analysis (using quantitative real-time reverse transcription polymerase chain reaction, qRT-PCR). Next, fetal lung progenitors (fLP) from stage 3 of the differentiation were generated by adding Wnt pathway activator CHIR99021 and retinoic acid (both of which influence lung specification^31^) to media supplemented with fibroblast growth factors (FGFs) and BMP4 (favoring conditions that drive fetal lung growth^18^). Gene expression analysis by qRT-PCR of the cell cultures at this stage showed abundant levels of *CFTR* expression (**Supplementary Fig. 2**). This high expression level of *CFTR* is reminiscent of first trimester fetal lung tissue, in which developing epithelium abundantly expressed CFTR^32^. An increase in gene expression associated with basal (*TRP63*, transformation related protein 63 or TP63; and cytokeratin-5, *KRT5*) and ciliated (forkhead box J1, *FOXJ1*) cells was also observed.

**Fig 1.**
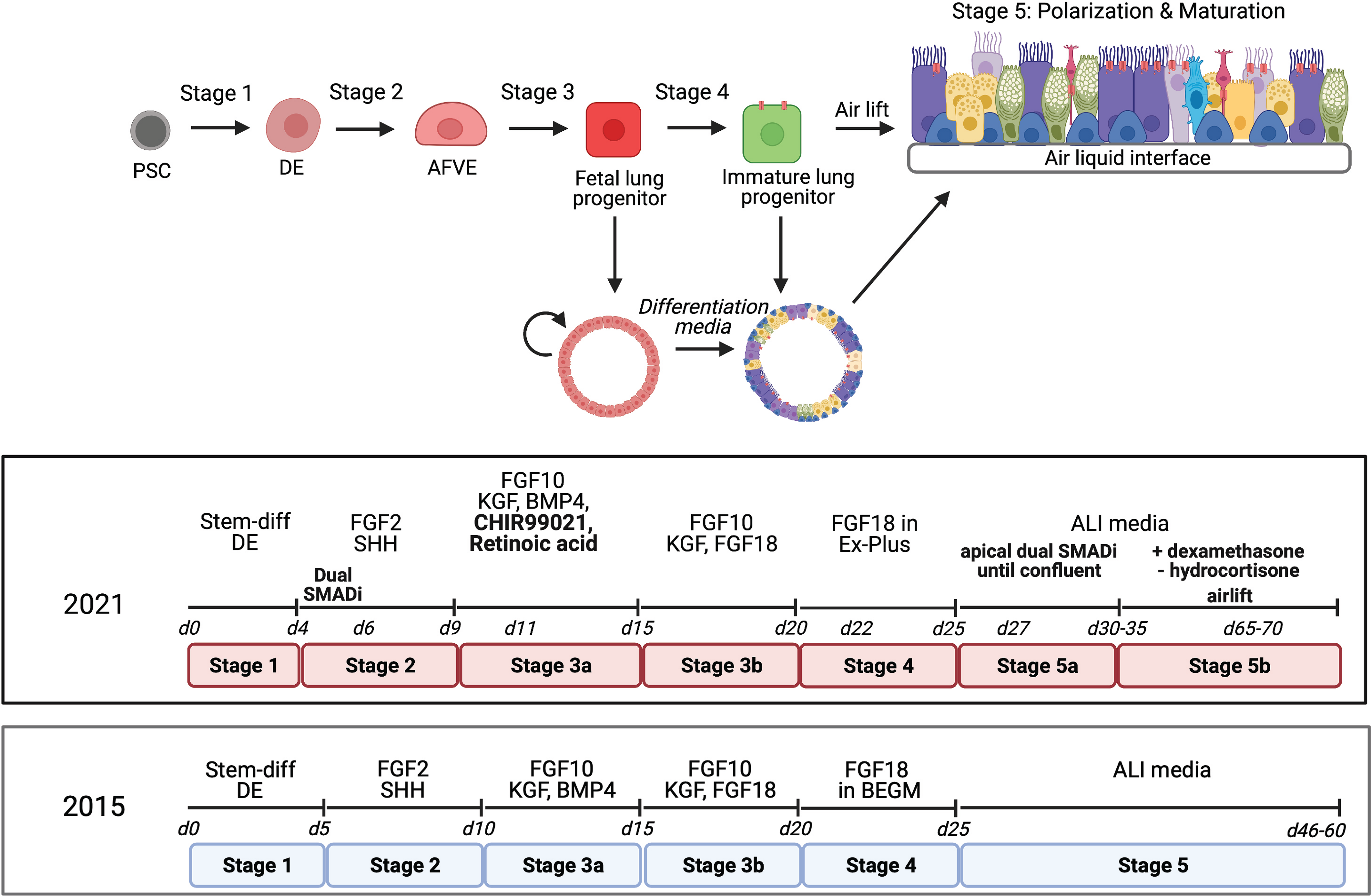
Refined *in-vitro* step-wise differentiation of human pluripotent stem cells to generate fetal lung (Stage 3) cells, immature spheroids (Stage 4) and mature ALI cultures (Stage 5). *Top*: Renewable fetal lung spheroids can be generated from hPSC-derived fetal lung (Stage 3) cells and further differentiated to generate immature lung spheroids (Stage 4) that can be dissociated onto transwells to culture in air liquid interface (ALI) for at least 5 weeks. *Bottom:* Detailed timeline comparing the refined differentiation method to our previously published method^22^.

To determine the heterogeneity of cell types and cell states in our *in-vitro* fLP culture, scRNA-seq was performed (**Fig. 2A**). We first compared fetal lung cells from our new differentiation to lung cells isolated from fetal lung tissue from two gestational week (GW) lungs based on a Pearson correlation coefficient (**Fig. 2B**). Our fetal lung cells had high correlation with both GW16 and GW18 lungs (> 0.85), however, most clusters in our culture had higher correlation to our GW16 lungs, with the exception of NKX2-1^hi^SOX9^hi^SOX2^lo^. Uniform manifold approximation and projection (UMAP) revealed the most common cell types found in both new and old differentiation fetal lung cell cultures, with the exception of a cluster of NKX2- 1^neg^SOX9^lo^SOX2^hi^ in our old protocol that lack *NKX2-1* expression but share similar expression profile as NKX2-1^hi^SOX9^hi^SOX2^lo^ cluster (**Fig. 2C**). First, all “lung progenitor” cells expressed high levels of pan-epithelial genes *EPCAM* and *CDH1* (E-cadherin), as well as *ID1* (Inhibitor of DNA binding 1) (**Fig. 2D**, **Supplementary Fig. 3A**). Within the lung progenitor population, the clusters are subdivided by their expression of *NKX2-1, SOX9* and *SOX2* which is previously suggested to be NKX2-1^hi^SOX9^hi^SOX2^lo^(bud tip), NKX2-1^hi^SOX9^lo^SOX2^hi^ (stalk), and NKX2- 1^hi^SOX9^lo^SOX2^lo^ (potentially developing saccule) progenitor cells^31^. Amongst the subdivided populations, the NKX2-1^hi^SOX9^lo^SOX2^lo^ lung progenitor cluster expressed elevated levels of sonic hedgehog signaling molecule (*SHH*) and *SOX17* which are associated with branching morphogenesis during fetal lung development^33, 34^, and high level of *CFTR* which may play a role in branching regulation^35^. The cluster of NKX2-1^hi^SOX9^hi^SOX2^lo^ lung progenitors expressed even higher levels of *ID1*, *ID2*, *ID3*, as well as Forkhead Box P1 (*FOXP1)*, Gata-binding protein-6 (*GATA6*), and Forkhead Box A2 (*FOXA2),* reminiscent of embryonic tip cells^31^. A cluster of NKX2-1^hi^SOX9^lo^SOX2^hi^ lung progenitors expressed high levels of cytokeratin-7 (*KRT7)*, insulin-like growth factor binding protein-5 (*IGFBP5*), and long non-coding RNA that harbours the gene miR-205 (*MIR205HG,* expressed in epithelial cells), reminiscent of embryonic stalk cells^31^. Specialized cell clusters, including basal, ciliated, pulmonary neuroendocrine cells (PNEC), and mesenchymal cells, were also found for both protocols. Basal cells expressed the canonical marker *TP63* as well as *ID1, KRT19,* TNF receptor superfamily member 12A (*TNFRSF12A*), and *MIR205HG*, while both cycling and non-cycling ciliated cells expressed *FOXJ1* and centromere protein U (*CENPU*)^36^. Cycling ciliated cells also expressed high levels of the proliferation gene DNA topoisomerase II alpha (*TOP2A*) while ciliated cell precursors expressed high levels of *ID1*. Ciliated cells expressed high levels of Forkhead Box N4 *(FOXN4*). Pulmonary neuroendocrine cells (PNEC) expressed the canonical gene achaete-scute family BHLH transcription factor 1 (*ASCL1*), as well as chromogranin B (*CHGB*), delta like canonical notch ligand 3 (*DLL3*), and myocardial infarction associated transcript (*MIAT*) while secretory precursors expressed secretoglobulin 3A2 (*SCGB3A2*), *ID1,* marker of proliferation Ki-67 (*MKI67*)*, TOP2A,* ADAM metallopeptidase domain 28 (*ADAM28*), and *MIR205HG*. Within the mesenchymal clusters, PDGFRB^hi^ cells expressed the highest level of platelet-derived growth factor receptor-b (*PDGFRB)*, as well as collagen alpha-1 (*COL1A1*) and *TOP2A*, suggesting these are cycling mesenchymal cells. On the contrary, PLAT^hi^ mesenchymal cells expressed high levels of plasminogen activator, tissue type (*PLAT*), and lower *COL1A1*.

**Fig 2.**
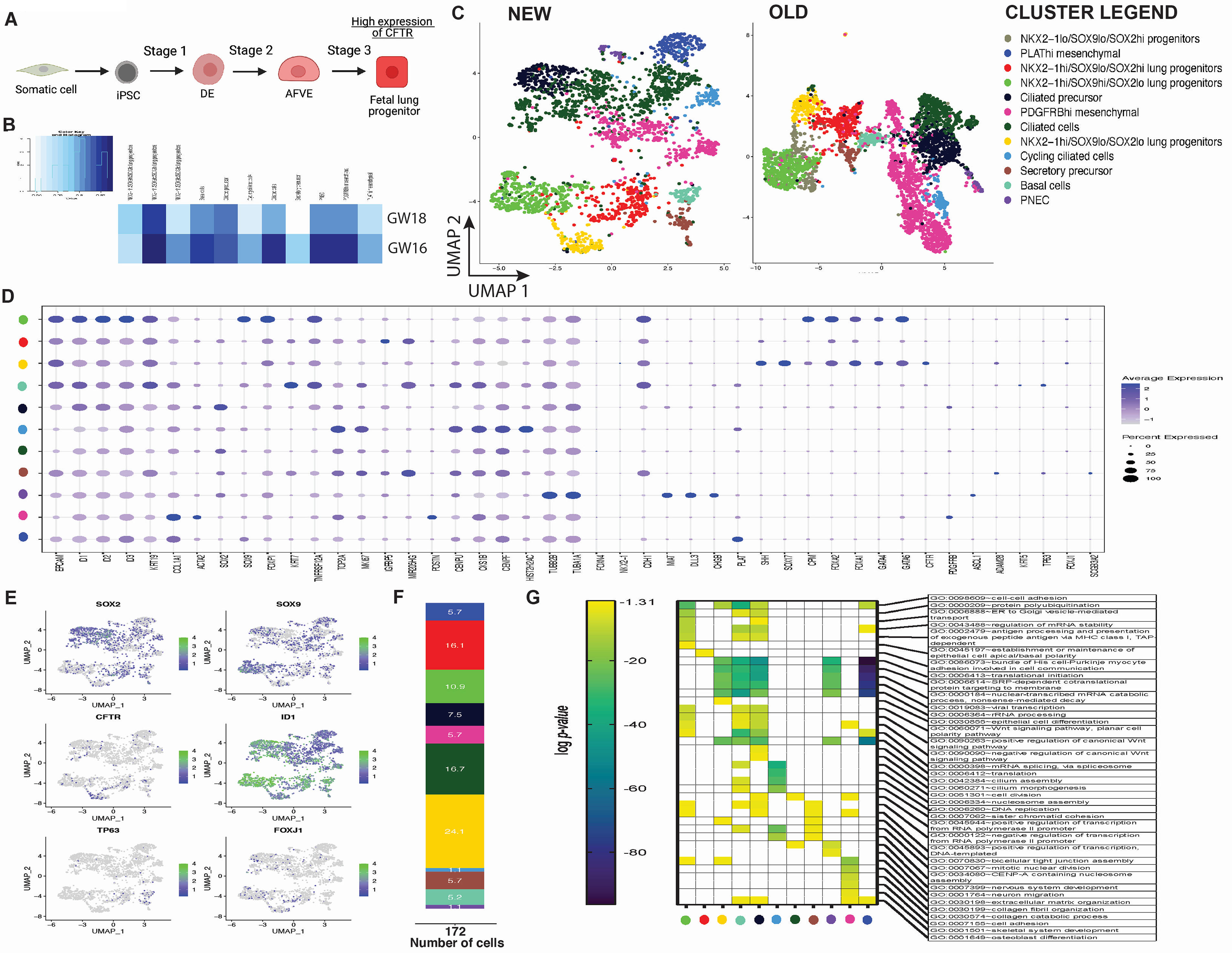
Stage-matching *in-vitro* differentiated hPSC-derived fetal lung (fLP) cells to *in-vivo* human fetal lung tissue. *A*. Differentiation to stage 3 fetal lung (fLP) cells. ***B*.** Pearson correlation analysis of upregulated genes between hPSC-derived fetal lung cells and primary fetal samples from pseudoglandular (GW16) and early-canalicular (GW18) shows hPSC-derived fLP are highly correlated with pseudoglandular fetal lung tissues (GW16) and early canalicular fetal lung tissues (GW18) (p >0.85). Reference map shows the Pearson correlation coefficient range from 0.855 to 0.924. ***C.*** Single cell RNA sequencing of *in-vitro* differentiated fLP for both new and old protocols. A total of ∼3000 cells (per protocol) were analyzed, and the single cell transcriptome was visualized by the UMAP plots of unsupervised clustering of cells for both new and old protocols. ***D.*** Average expression level for each gene and frequency of cells expressing the gene is shown by a dotplot of fLP derived from the new protocol. Reference map shows intensity of the average gene expression level and percentage of specific gene-expressing cells are demarcated by size of the circles. ***E.*** Feature plots of select gene expression show distribution of the expression in the UMAP clusters. Reference bar represents the avg_log2FC. Minimum value set as 0.2. ***F.*** The frequency of CFTR-expressing cells in all clusters. ***G.*** Heatmap represents the top (up to) 5 and cluster-specific gene ontology (GO) terms identified with each cluster based on differential gene expression (DEG). The Benjamini-adjusted p-value in log scale (<0.05 or log (-1.3)) was used to determine DEG. Empty boxes represent no or insignificant p-values. Note: each cluster is colour coded based on the UMAP clusters in **C**.

Feature map of the genes showed that *SOX2* was predominantly expressed in NKX2- 1^hi^SOX9^lo^SOX2^hi^ progenitor cells, ciliated cell populations (ciliated cells, ciliated precursor, cycling ciliated cells), and PNEC, suggesting proximal lineages. *SOX9*, however, is expressed predominantly in NKX2-1^hi^SOX9^hi^SOX2^lo^ and sporadically in PDGFRB^hi^ mesenchymal cells and NKX2-1^hi^SOX9^lo^SOX2^hi^ clusters, indicative of distal lineages (**Fig. 2E**). *ID1* is highly expressed in ciliated precursors and all lung progenitor cells. *FOXJ1* and *TP63* are expressed in ciliated cell and basal cell clusters respectively. We also found *CFTR* expressing cells predominantly in the lung progenitor populations of NKX2-1^hi^SOX9^hi^SOX2^lo^ (10.9%), NKX2-1^hi^SOX9^lo^SOX2^lo^ (24.1%), NKX2-1^hi^SOX9^lo^SOX2^hi^ (16.1%), and ciliated cells (16.7%) (**Fig. 2F**).

Next, we assessed the Gene Ontology (GO) terms associated with each cluster based on the differentially expressed genes (DEG) in both protocols and found processes related to cell signaling, proliferation, adhesion, motility, transcription and developmental processes (**Fig. 2G** and **Supplementary Fig. 3B**). Among the top 5 most significant GO terms in each cluster, we found all epithelial cell clusters (all lung progenitors, basal cells, all ciliated cells, secretory precursors and PNEC) and PLAT^hi^ mesenchymal cells expressed genes related to biological pathways concerning protein synthesis (e.g. regulation of mRNA stability, translational initiation, protein polyubiquitination). Additionally, GO terms related to cell division (cell division, mitotic nuclear division, sister chromatid cohesion) had significant presence in basal cells, cycling ciliated cells, ciliated cells, and secretory precursors, supporting the proliferative capacities of these cell types during the fLP stage. Interestingly, NKX2-1^hi^SOX9^lo^SOX2^lo^, NKX2-1^hi^SOX9^hi^SOX2^lo^, and NKX2-1^hi^SOX9^lo^SOX2^hi^ lung progenitors contain significant unique GO terms associated with epithelial cell differentiation, establishment or maintenance of epithelial cell apical/basal polarity, and muscle-interacting biological pathways respectively. The unique GO terms among the three progenitor clusters resemble the different roles of lung progenitors during branching morphogenesis in late pseudoglandular stage^31^. PNEC are significantly associated with multiple neurological-related pathways (e.g. sympathetic nervous system development, neuron migration), which supports their role in neurological sensory function^37^. Ciliated cells associated with GO terms relevant to positive regulation of transcription from the RNA polymerase II promoter and nervous system development, while cycling ciliated cells and ciliated precursors were associated with biological processes of cell proliferation and transcription-related pathways respectively. Importantly, ciliated precursors were significantly associated with processes such as cilium assembly and cilium morphogenesis. Basal cells and ciliated precursors also expressed genes associated with Wnt signaling pathways (e.g. Wnt signaling pathway, planar cell polarity pathway, positive/negative regulation of canonical Wnt signaling pathway) which were previously shown to regulate ciliated cell differentiation^38^. PLAT^hi^mesenchymal cell GO analysis revealed processes related to protein synthesis pathways and non-lung differentiation pathways. GO analysis of PDGFRB^hi^ mesenchymal cells, on the other hand, indicated processes related to extracellular matrix, bone, and muscle. The distinctive GO terms between PLAT^hi^ and PDGFRB^hi^ mesenchymal cells suggest different phenotype and functional roles for these cells in the fLP cultures.

Immunofluorescence staining of fLP cultures for multipotent progenitor cell markers (SOX2, SOX9) and lung marker NKX2-1 (or thyroid transcriptional factor 1, TTF1, protein *herein*) showed TTF1^+^ cells expressing both SOX2^+^ and SOX9^+^ as well as TTF1^+^SOX2^+^ and TTF1^+^SOX9^+^ double positive cells were found for both protocols (**Fig. 3A** and **Supplementary Fig. 3C**) using both hPSC lines, suggesting emergence of a heterogeneous lung progenitor population in the fLP cultures. Using fluorescence activated cell sorting (FACS) analysis, a general increase in TTF1+ cells as well as TTF1^+^SOX2^+^ and TTF1^+^SOX9^+^ double positive cells was observed (**Fig. 3B**). This is in line with findings pertaining to multipotent progenitors that express both SOX2 and SOX9 in the distal tip of the developing human lung^31^. It is also in line with previous differentiation studies demonstrating fetal-like lung progenitors after differentiating anterior ventral foregut endoderm cells.

**Fig 3.**
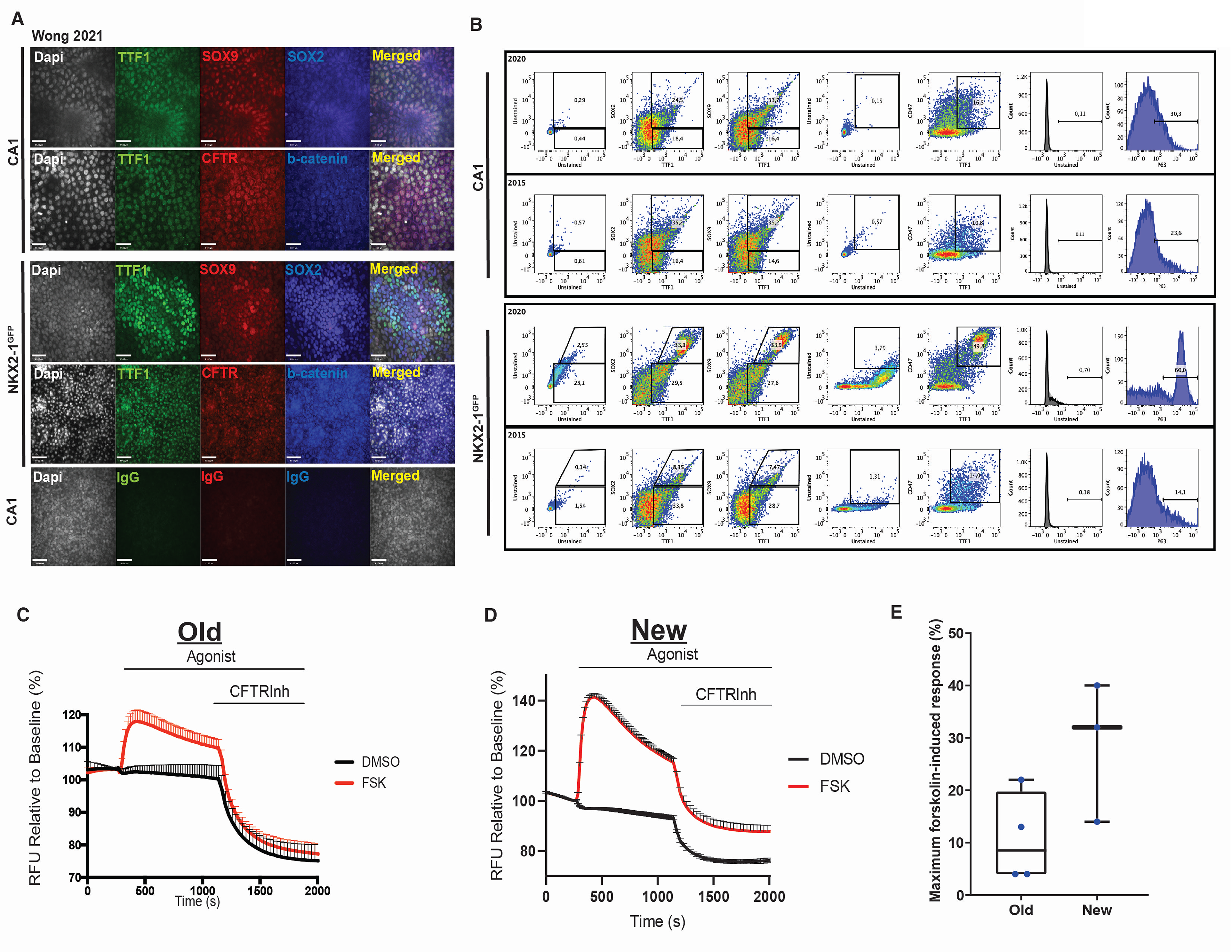
Phenotype and functional validation of *in-vitro* differentiated fetal lung progenitors (fLP). ***A.*** Immunofluorescence staining of fLP for proteins highly expressed in the developing fetal lung tissue TTF1, SOX2, SOX9, CFTR and β-catenin. Two human pluripotent stem cell lines (CA1 and BU3-NKX2-1^GFP^) were used. Non-immune isotype controls were used to determine non-specific binding of secondary antibodies. White bar represents 41 microns (low magnification) and 20 microns (high magnification) respectively. ***B.*** Flow cytometric analysis demonstrating generation of lung (TTF1, CD47) cells expressing the basal stem cell marker (ΔNP63), proximal (SOX2) and distal (SOX9) in the new and old protocol. ***C, D***. Representative apical chloride conductance traces of fLP demonstrate robust CFTR functional response to forskolin-induced channel activity (red) that is inhibited by CFTR inhibitor-172 (n=5) in old and new protocols, respectively. ***E.*** Representative Western Blot images highlighting mature (Band C) and immature (Band B) CFTR protein in old and new protocols. WT and KO-HBE cells were used as positive and negative controls respectively. Calnexin was used as a loading control. ***F.*** Quantitative box-plots comparing maximal forskolin-induced CFTR activation (relative to unstimulated baseline reads) in independent sets of fLP from the old and new differentiation protocols.

It has previously been shown that an integrin associated protein, CD47, can be used to isolate NKX2-1^+^ progenitors^39^. Similarly, in our fLP cultures, a significant increase in CD47^+^TTF1^+^ double positive cells were found. Moreover, fLP from the new differentiation protocol expressed substantially more plasma membrane-localized CFTR compared to our previous protocol (**Fig. 3A**).

Recent studies have identified fetal basal stem cells at the fLP stage of differentiation as marked by expression of *TP63*^11, 13^. We also showed improved generation of P63^+^ basal stem cells (>16% in CA1 and >51% in BU3-NKX2-1^GFP^) in our new protocol, and found differences in efficiency in two independent PSC lines (**Fig. 3B**). Indeed, we observed increased expression levels of genes associated with basal cells (*TRP63* and *KRT5*) as well as ciliated cells (*FOXJ1*) (**Supplementary Fig. 2**).

Using our previously established modified apical chloride conductance (ACC) assay to measure membrane potential changes through changes in fluorescence intensity (via a fluorescence imaging plate reader, FLiPR^23^), we found CFTR activity was enhanced by approximately 30% in our new protocol compared to our old (**Fig. 3C,D)** upon forskolin-stimulation. Specificity of CFTR function was confirmed using CFTR inhibitor-172 treatment. Higher protein levels of the mature CFTR protein Band C (representing a fully glycosylated functional form of the protein expressed on the plasma membrane, of a size of ∼170 kDa) was observed in our new differentiation protocol compared to our previous protocol (**Fig. 3E**). Both hPSC lines demonstrated enhanced CFTR function averaging around 32% function above baseline prior to forskolin-stimulation, in our new protocol compared to our old protocol, which produced ∼10% function (**Fig. 3F**). As such, this provides a powerful source of cells for rapid high throughput functional screens of novel CFTR modulators^28^.

### Generation of renewable 3D lung spheroids

To determine whether isolated fLP embedded in matrigel could generate lung epithelial spheroids (**Fig. 4A**), we sorted for pan-epithelial cell adhesion molecules (EPCAM or CD326) and found the majority of the fLP were CD326^+^ (up to 72%, **Fig. 4B**), now called fetal lung epithelial cells (fLECs). Within 4 days, epithelial spheroids emerged (**Fig. 4C**). These spheroids were capable of propagation in expansion media (**Fig. 4D**) containing dual smad inhibitors for up to 10-15 passages with decreasing expansion potential in late passages (data not shown). We tested this spheroid generation potential in three independent hPSC lines and found varying efficiencies of spheroid generation (**Fig. 4E, F**) from fLECs. Renewable epithelial spheroids are composed of mostly ΔNP63^+^ basal stem cells (>90%, **Fig. 4G, H**), confirmed using a double fluorescence reporter hPSC line BU3-NKX2-1^GFP^/P63^tdTomato^ (NGPT)^13^ where tdTomato fluorescence was observed in the majority of the cells in the spheroids. The abundance of basal stem cells suggests the expansion capacity of the spheroids may be driven by these multipotent progenitor cells as previously described^13^. The spheroids can be cryopreserved and expanded without losing cell phenotype after serial passaging (data not shown). Phenotypic characterization of the self-renewing spheroids shows cells expressing TTF1, SOX2, SOX9 and luminal expression of tight junction protein (ZO- 1). CFTR appears localized to the cytoplasm of the cells (**Fig. 4I**). Notably, the majority of these cells express ΔNP63 and cytokeratin-14 (KRT14), associated with basement membrane localized basal cells in the airways.

**Fig 4.**
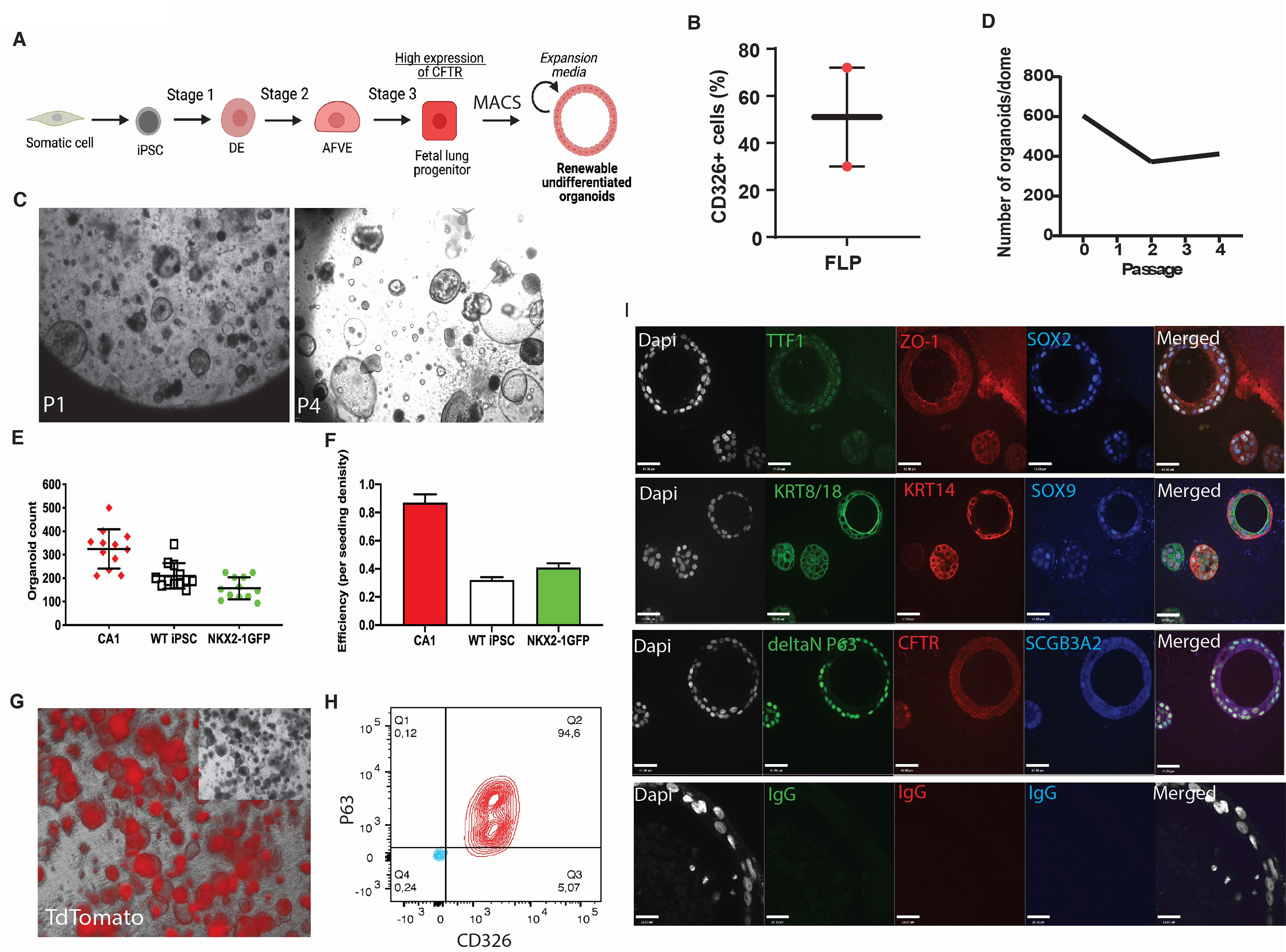
Phenotype and expansion characterization of *in-vitro* proliferative lung epithelial spheroids. *A*. Differentiation to fLEC airway epithelial progenitor spheroids. ***B*.** Percentage of CD326^+^ expressing epithelial cells in fetal lung cells assessed by magnetic-activated cell sorting assay. ***C.*** Representative DIC image of airway epithelial spheroids at passage 1 and 4. ***D.*** Expansion curve of spheroids of number organoids per dome. ***E.*** Number of epithelial spheroids generated from 3 different hPSC lines. ***F*.** Efficiency of generating epithelial spheroids range from 30-85% depending on the line. ***G.*** Representative images of BU3-NKX2-1^GFP^/P63^dTomato^ in DIC and immunofluorescence imaging for P63^tdTomato^ reporter*. **H.*** Flow cytometry analysis showed high levels of basal (ΔNP63) and epithelial (CD326) in stage 3 spheroids. Red and blue contour plots represent stained and unstained cells, respectively. ***I.*** Immunofluorescence images shows undifferentiated lung epithelial spheroids expresses fetal lung (TTF1, SOX2, SOX9), epithelial (KRT8/18), epithelial tight junctions (ZO-1) and pulmonary cell types (ΔNP63, Basal stem cell; KRT14, Basal cells; SCGB3A2, secretory cells). Rabbit primary antibodies shown in green, mouse primary antibodies in red and goat or rat primary antibodies shown in blue. Non-immune reactive isotypes (IgG) were used as control staining. White scale bar is automatically generated and represents 41 microns (low magnification) and 20 microns (high magnification) images.

To determine the multi-epithelial differentiation potential of the basal stem cells in the epithelial spheroids, the spheroids were cultured in differentiation media for up to 14 days (**Fig. 5A**). Morphologically, hPSC-derived epithelial spheroids share morphological similarities to spheroids derived from primary bronchial epithelial cells (**Fig. 5B**). Characterization of the differentiated spheroids show up-regulation of several genes associated with specific epithelial cell lineages found in the airways including basal cell marker KRT5, ciliated cell marker FOXJ1, luminal epithelial cell marker KRT8, and secretory cell marker MUC16 (**Fig. 5C**) and is confirmed with the presence of ΔNP63 and KRT14, acetylated a-tubulin, cytokeratin-8/18 (KRT8/18), and SCGB1A1 and mucin 5ac (MUC5AC) respectively in bulk qPCR analysis (**Fig. 5D**). A near two-fold up-regulation of *CFTR* is also observed and can be found on the apical membrane (lumen side) of the spheroid along ZO-1. The lung (TTF1^+^) epithelial (E-cadherin^+^, panCK^+^) spheroids express proximal marker SOX2 suggesting these are airway epithelial spheroids and do not contain cells of the distal respiratory phenotype.

**Fig 5.**
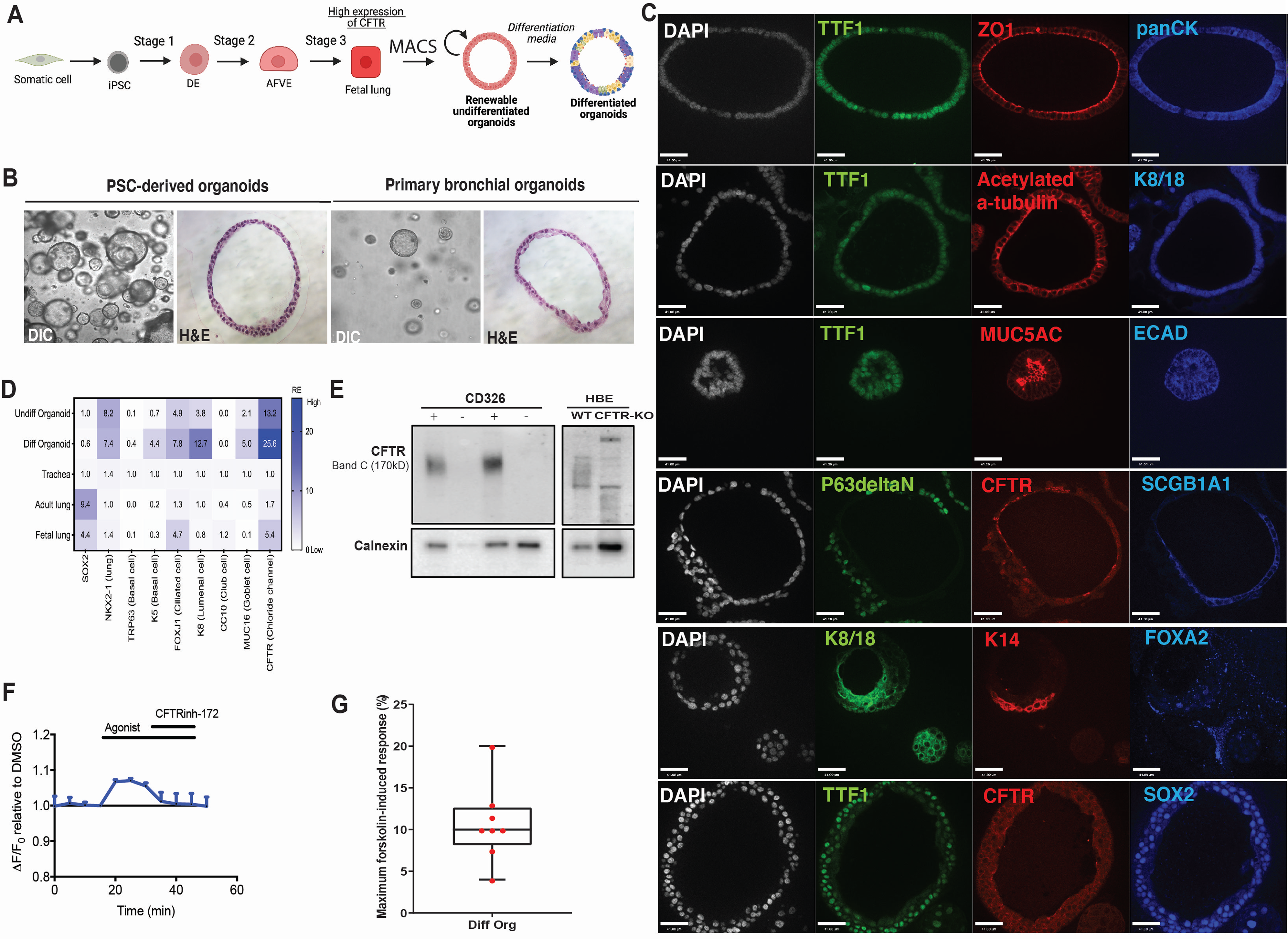
Phenotype and functional characterization of *in-vitro* differentiated lung epithelial spheroids. ***A.*** Differentiation of airway epithelial spheroids. ***B.*** Representative DIC and H&E images of *in-vitro-*derived airway epithelial spheroids and primary bronchial cells-derived airway spheroids. ***C*.** Characterization of *in-vitro*-derived airway epithelial spheroids for lung airway (SOX2, TTF1), pan-epithelial (panCK, E-cadherin) and specific epithelial cell markers (ZO1, tight junction; acetylated alpha tubulin, cilia on ciliated cells; MUC5AC, goblet cell; ΔNP63, basal stem cell; KRT14, basal cell; SCGB1A1, club cell). Epithelial spheroids express mature luminal epithelial markers KRT8/18 and luminal expression of chloride channel CFTR. Notably, pan-endoderm transcription factor FOXA2 is weakly expressed in a subset of cells in the spheroids. ***D.*** Heatmap of gene expression analysis of epithelial spheroids in undifferentiated (stage 3 organoids) and differentiated (stage 4 organoids) show an increase in expression of epithelial-specific lineage markers (*TRP63, K5, FOXJ1, K8, MUC16*) and a near 2-fold increase in *CFTR* expression. Genes were normalized to housekeeping gene *GAPDH* and expressed relative to RNA from trachea or adult lung (expression level = 1.0). At least 3 technical replicates were performed, n= 3 biological cell lines. ***E*.** Only epithelial (CD326^+^) cells (sorted by magnetic activated cell sorting (MACS) formed spheroids expressing the mature CFTR protein (Band C). Anti-calnexin was used as a control for protein loading. ***F*.** Representative ACC trace of CFTR function upon stimulation with forskolin and inhibited with CFTR inhibitor-172. ***G.*** Box-plot demonstrates the average maximum forskolin-induced response level of differentiated spheroids from different sets of spheroids. ***H.*** Representative brightfield images of spheroids pre (T0) and post-FIS at 24 hours (T24). White scale bar is automatically generated and represents 41 microns. ***I.*** Average diameter size of the spheroids at time 0 (T0) and 24 hours (T24) post FIS.

To assess CFTR protein expression, we assayed for the presence of Band C (∼170 kDa) protein by immunoblotting, as it is indicative of a mature apically localized protein capable of function. Only spheroids derived from CD326^+^ cells express mature CFTR (**Fig. 5E**). To validate this, we performed FLiPR analysis on spheroids 2-3 days^40^ after seeding on collagen-coated plates and found an increase in fluorescence activity indicative of CFTR function upon forskolin- induction that was inhibited by CFTR inhibitor-172 (**Fig. 5F**). Assessing the CFTR functional potential of several spheroids from different sets of differentiations shows, on average, up to 10% CFTR function can be detected in the differentiated spheroids (**Fig. 5G**). Importantly, a decrease in CFTR function in differentiated cells aligns with a decrease in CFTR expression in fetal lung tissues during development beyond the second trimester where CFTR is found only in a subset of bronchial epithelial cells^32, 41^. Since CFTR is expressed on the lumenal membrane of the spheroid, we also determined CFTR function by way of forskolin-induced swelling (FIS), as previously described in rectal organoids^42^. Spheroids were stimulated with 10µM of forskolin for 1, 6, and 24 hours. Modest swelling was observed between 1-6 hours (data not shown) but significant swelling was found after 24 hours of forskolin stimulation. Most spheroids at 24 hours appeared to have cystic-like morphology suggesting an over-extension of the epithelium (**Fig. 5H**). Quantification of the organoid size by diameter after 24 hours of FIS showed significant increase in size ranging from 250 µm to 600 µm, compared to 65-125 µm observed pre-FIS (P<0.0001, **Fig. 5I**).

### Generation of mature adult-like airway epithelial tissue

Next, we assessed the ability to generate functional mature airway epithelial tissue by culturing the cells on collagen-coated transwells (as previously described^18^) to produce air liquid interface (ALI) cultures (**Fig. 6A**). During this period, the epithelial cells undergo additional differentiation, maturation and polarization to form a pseudostratified epithelium reminiscent of the mature airways. Our modified ALI protocol entails a two-step method that first allows expansion of the cells seeded onto the transwell membrane to form a monolayer of cells prior to differentiation and polarization. Characterization of the genes expressed in the cell population of ALI cultures showed an average but significant up-regulation of *CFTR* expression in the new protocol by >500-fold compared to the old protocol, while many of the other epithelial cell lineage markers remained relatively consistent (**Supplementary Fig. 4)**. Importantly, there were differences in expression levels for many of the genes between the two hPSC lines tested.

**Fig 6.**
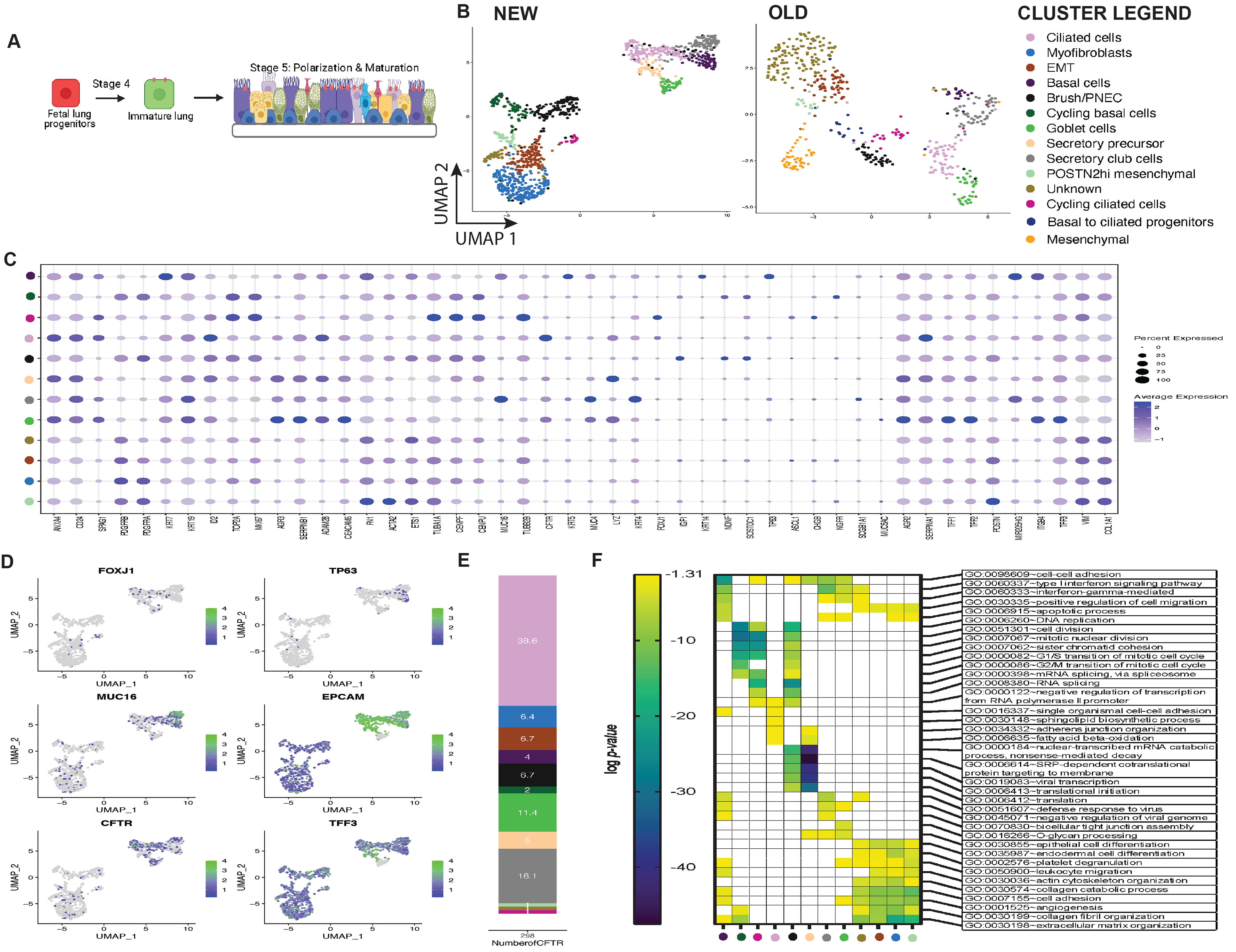
Single cell profiling of *in-vitro*-derived mature ALI airway cells compared to human bronchial epithelial cultures. *A*. Differentiation to mature airway epithelial cells in ALI for 5 weeks demonstrate. ***B***. A total of *in-vitro* ∼3000 cells per protocol were analyzed and the single-cell transcriptome was visualized by UMAP plot. Unsupervised clustering reveals 11 and 12 cell clusters in the old and new differentiation protocols, respectively. ***C.*** Dot plot shows the average gene expression of pulmonary cell-specific genes in each cluster for ALI cultures derived from the new protocol. Reference map shows the range of gene expression level and the number of cells in each cluster is represented by circle size. ***D.*** UMAP presents the average gene expression of selected genes in each cell. Reference bar represents the avg_log2FC. Minimum value set as 0.2. ***E.*** Frequency of CFTR-expressing cells in each cluster of mature ALI airway cells*. **F.*** Heatmap represents the top (up to) 5 gene ontology (GO) terms identified with each cluster based on differential gene expression (DEG). The Benjamini-adjusted p-value in log scale (<0.05 or log (- 1.3)) was used to determine DEG. Empty boxes represent no or insignificant p-values. Note: each cluster is colour coded based on the UMAP clusters in **B**.

For higher resolution of the cell heterogeneity in our cultures, we performed scRNAseq on the ALI epithelium after 5 weeks in culture. UMAP analysis displayed presence of additional cell types observed in the adult airways, including goblet cells, secretory club cells, ionocytes, and brush/PNEC cell types as well as cycling basal and ciliated cells (**Fig. 6B**), with the latter two expressing high levels of proliferation markers *MKI67* and *TOP2A* (**Fig. 6C** and **Supplementary Fig. 5A**)^5, 8, 10^. Basal cells expressed high levels of *TP63,* and keratins *KRT5, KRT14, KRT19,* and *KRT7* as previously described^5, 11^. Cycling basal cells expressed less *TP63* but higher sclerostin domain containing 1 (*SOSTDC1*), previously found in fetal basal cells with multipotent differentiation capacity^9^. Both cycling ciliated cells and ciliated cells expressed canonical marker *FOXJ1*. However, CFTR was predominantly expressed in the non-cycling ciliated cell population. Goblet cells expressed high levels of anterior gradient protein homologs *AGR2, AGR3,* lysozyme *LYZ,* trefoil factors *TFF1*, *TFF2, TFF3*. Secretory precursors expressed less secretory proteins and higher levels of *MKI67* and *TOP2A.* Secretory club cells expressed high levels of cytokeratins *KRT4, KRT19,* mucin *MUC16*, and secretoglobin *SCGB3A2*, while brush/PNEC expressed insulin growth factor *IGF1*, sclerostin domain containing protein *SOSTDC1*, and neuron derived neurotrophic factor *NDNF*. A cluster of unknown cells expressing high levels of IQ motif containing GTPase activating protein 2 (*IQGAP2*)*, ID3,* and *ID4* may be pre-ionocyte^8^ cells, although without expression of the canonical genes *FOXI1* and *CFTR,* the phenotype remains unresolved. Interestingly, *CFTR* was expressed in most epithelial cell types including goblet, secretory club, ciliated, basal cells and unlike previous findings^8^, to a lesser extent in the unknown cell type. Epithelial cells undergoing epithelial-mesenchymal transition (EMT) were also found in the ALI cultures and expressed both epithelial cytokeratins (*KRT19, KRT7*) and canonical mesenchymal cell genes platelet-derived growth factor receptors *PDGFRA*, and *PDGFRB*, suggesting the need for supporting mesenchymal cells in the maturation of the airways^8, 43^. Our new differentiation protocol also identified a cluster of POSTN2^hi^ mesenchymal cells that express genes commonly found in the myofibroblast cluster with the exception of a slightly higher expression of Periostin (*POSTN2*), previously found in human lungs that secrete extracellular matrices such as collagen^44^. In our model, *CFTR* expression is highly expressed within the ciliated cell cluster (**Fig. 6D**). While an unknown cluster of cells that may represent pre-ionocyte cells exist, these cells do not express *CFTR*. Assessing the number of CFTR-expressing cells in the ALI culture shows the majority of these cells are ciliated, followed by goblet and secretory club cells (**Fig. 6E**). The reduced expression of *CFTR* in clusters associated with EMT and myofibroblasts implicates a role of CFTR in epithelial-mesenchymal transition^2^.

Gene ontology analysis of each cluster showed the top 5 most significant GO terms were generally related to cell cycle regulation, cell matrix organizations, immune pathways and developmental signaling pathways for the ALI cultures (**Fig. 6F** and **Supplementary Fig. 5B**). We found all clusters in both protocols contained GO terms related to multiple cell adhesion/matrices-related biology pathways (cell-cell adhesion, adherens junction organization, cell adhesion, bicellular tight junction assembly, collagen catabolic process) which resembles the active transition of extracellular matrices in the lung^45^. Cell division, mitotic nuclear division, sister chromatid cohesion, and G1/S transition of mitotic cell cycle were found in cycling basal cells, cycling ciliated cells and brush/PNEC, indicative of the proliferative nature of these cell types. We also found immune response related GO terms in secretory and basal cells which supports their role in regulating immunity in the epithelium^46^. Sphingolipid biosynthesis is important in regulating cilia functions and formations in ciliated cells^47^. Notably, the unknown cluster had GO terms associated with cell proliferation (positive/negative regulations of proliferation), extracellular matrix (extracellular matrix organization, collagen fibril organization) and angiogenesis-related (angiogenesis, blood vessel development, sprouting angiogenesis) biological pathways.

Immunostaining of the mature ALI cultures in our new protocol showed TTF1 expression was restricted to certain cell types, and the majority co-expressed with SOX2, indicating proximal airway phenotype (**Fig. 7A** and **7A’**), similar to adult lung tissue (**Fig. 7A”**). A polarized pseudostratified airway epithelium is formed after 5 weeks of ALI composed of basal cells (ΔNP63^+^) in the basolateral side of the culture, with secretory (SCGB3A2^+^), and ciliated (acetylated alpha-tubulin^+^) cells extended to the apical side (**Fig. 7B**), similar to adult airway tissue (**Fig. 7B’**). Non-immune reactive immunoglobulins (IgGs) were used as controls for non-specific binding (**Fig. 7C**). Bulk gene expression analysis showed both hPSC lines were capable of generating many of the mature airway epithelial cell types observed in airways using the new protocol (**Supplementary Fig. 4**). Of the FOXJ1-expressing cell population, 19% of cells co-expressed CFTR and FOXJ1, confirming that a subpopulation of ciliated cells express CFTR, while smaller percentages of ΔNP63+ basal cells and SCGB3A2+ secretory cells also expressed CFTR (**Fig. 7D**). Protein analysis of CFTR shows high levels of band C (∼170 kDa), representing the mature fully glycosylated form of the CFTR protein (**Fig. 7E**). Notably, a near 3-fold higher CFTR function was detected in the new ALI cultures compared to our previous protocol (**Fig. 7F, G**).

**Fig 7.**
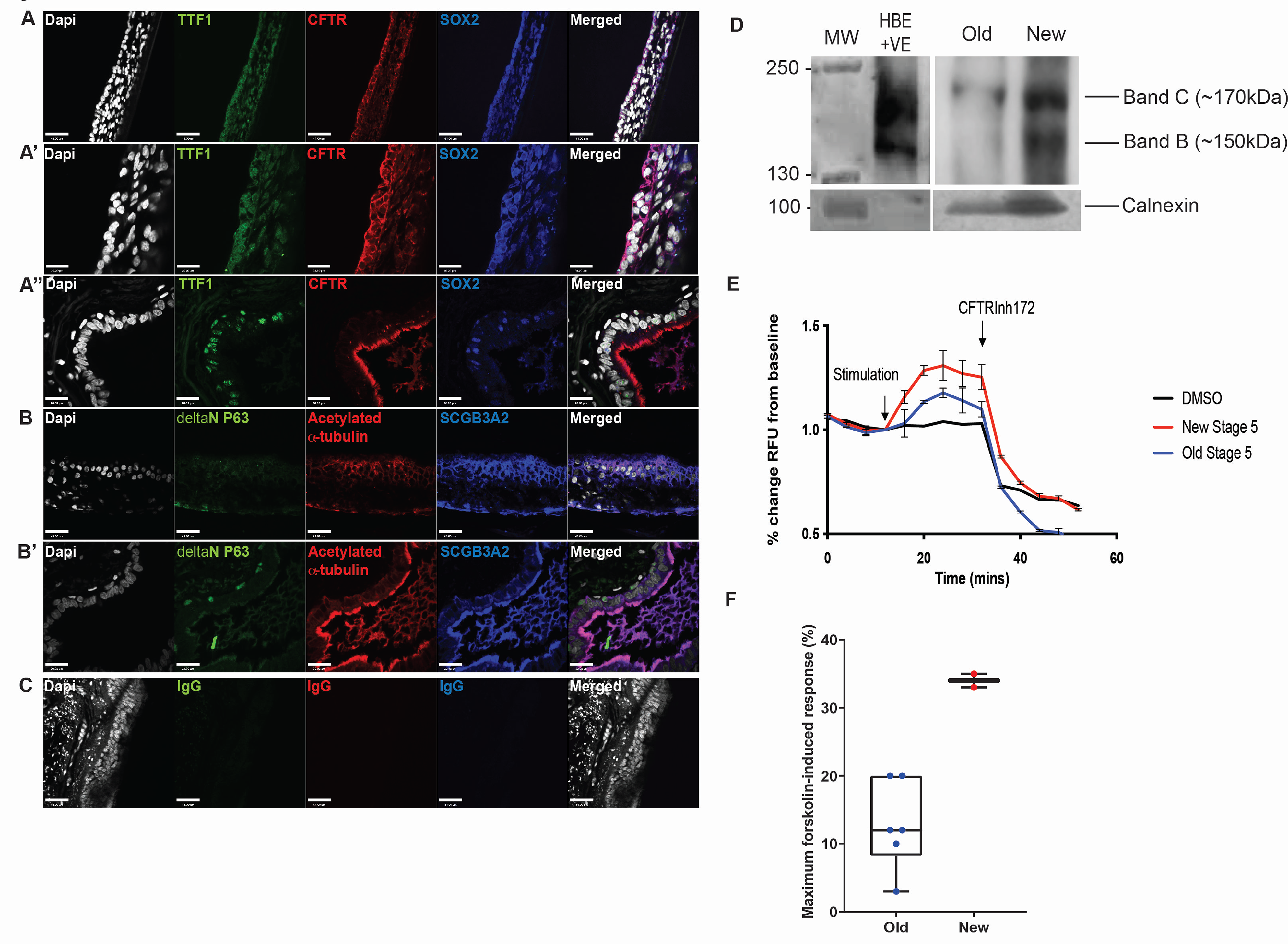
Phenotype and functional validation of *in-vitro* differentiated mature airway epithelial cells. ***A*** and ***A’*.** Immunofluorescence staining of ALI cultures (low and high magnification, respectively) from Stage 5 cells show proximal (SOX2) airway (TTF1) phenotype which is reminiscent of a pseudostratified structure in control lung tissue ***A’’***. ***B.*** Basal (ΔNP63), secretory (SCGB3A2) and ciliated (acetylated alpha-tubulin) cells are present and form a polarized epithelial similar to control primary adult lung airway and ***B’. C.*** Non-immune isotype controls were used to determine non-specific binding of secondary antibodies. White bar represents 41 microns and 20 microns for low and high magnification, respectively. ***D*.** Flow cytometric analysis for CFTR and various lung epithelial cell markers (FOXJ1 for ciliated cells, ΔNP63 for basal stem cells, and SCGB3A2 for secretory cells). ***E*.** Immunoblot analysis for CFTR protein species shows higher levels of a mature CFTR protein (Band C, ∼160kDa) in stage 5 cells in the new differentiation to mature airway epithelia derived from both human PSC lines. Anti-calnexin was stained to evaluate protein loading. Human bronchial epithelial cells (HBE WT-CFTR) expressing wild-type CFTR were used as positive control whereas CFTR knock-out in human bronchial epithelial cells (HBE KO-CFTR) was used as negative control. ***F.*** Representative apical chloride conductance trace of mature lung cells demonstrating robust CFTR functional response to forskolin-induced channel activity (red) that is inhibited by CFTR inhibitor-172 (n=5). ***G*.** Comparing maximal forskolin- induced CFTR activation of stage 5 cells demonstrates a 2-fold higher level of functional activity in the new differentiation protocol.

### Cell lineage relationship of differentiated cells

Cells that differentiate display a continuous spectrum of states simply because each cell will differentiate in an unsynchronized manner (i.e. each cell represents a snapshot of differentiation in time). Therefore, the distance between a cell and the start of a differentiation trajectory (or pseudotime) can be inferred. Here we used 2 known methods of trajectory inference: Monocle 3 pseudotime and scVelo RNA velocity. First, to determine the lineage relationships of the cells as they differentiate from fetal to ALI stages, we performed pseudotime analysis of the integrated fLP and ALI clusters using Monocle3^48^. To do this, we combined the data sets of fLP and ALI cultures using Seurat V3’s built-in integration algorithm^49^ which identified anchor genes between the two data sets that were used to subsequently merge the datasets. After integration, we first generated an integrated UMAP outlining the cell clusters (**Fig. 8A**). Pseudotime analysis was then performed to infer the cell lineage trajectories and the temporal relationship of the cells based on gene expression changes through pseudotime. Using the built-in partition-based graph abstraction algorithm in Monocle3^50^, the integrated clusters were partitioned into 5 groups (‘parts’) (**Fig. 8B**), and within each ‘part’ cell trajectories were identified (**Fig. 8C**). A limitation of using Monocle3 pseudotime analysis is that the trajectory is based on a user-defined starting node. We also leveraged RNA abundance (RNA velocity) as a measure of cell state and determined developmental relationships and cellular dynamics for each cell^51^ (**Fig. 8D**). Based on both pseudotime and RNA velocity, we found that Part 1 and 4 appeared to share similar lineage trajectories. First, a relationship between ciliated cells and brush/PNEC and PLAT^hi^ mesenchymal cells was revealed. Myofibroblasts (ALI) emerged from several cell types: ciliated cells and various mesenchymal populations (PLAT^hi^, POSTN2^hi^ (ALI) and PDGFRB^hi^ mesenchymal cells). In part 2, pseudotime analysis revealed two distinct lineage trajectories. First, NKX2- 1^hi^SOX9^lo^SOX2^hi^ and NKX2-1^hi^SOX9^hi^SOX2^lo^ progenitor cells appear to have interchangeable flow velocity trajectories. NKX2-1^hi^SOX9^lo^SOX2^lo^ progenitors emerge from NKX2- 1^hi^SOX9^hi^SOX2^lo^ but are largely cycling. We observed a relationship between NKX2- 1^hi^SOX9^lo^SOX2^lo^ progenitors and secretory precursors and goblet cells. A small subset of fetal lung ciliated cells and secretory precursors share RNA velocity trajectories with cycling ciliated cells. Also discovered was a lineage relationship between basal cells in the fetal stage with basal cells in ALI and secretory club cells and the latter with ciliated cells. Finally, a lineage relationship between fetal lung ciliated cells with NKX2-1^hi^SOX9^hi^SOX2^lo^ progenitors and secretory precursor cells was also found. In part 3, a relationship between fetal lung ciliated cells and ciliated cell precursors was found, as well as cycling basal cells with ciliated cell precursors. Interestingly, there appears to be overlap between cycling ciliated cells in ALI with ciliated cells in the fetal lung stage, suggesting a potential “de-differentiation” or reversion to a premature state during cell proliferation. Finally, the last partition was largely dominated by fetal lung PNEC, suggestive of an independent cell cluster that has no obvious lineage relationships to other cell types found in the culture.

**Fig 8.**
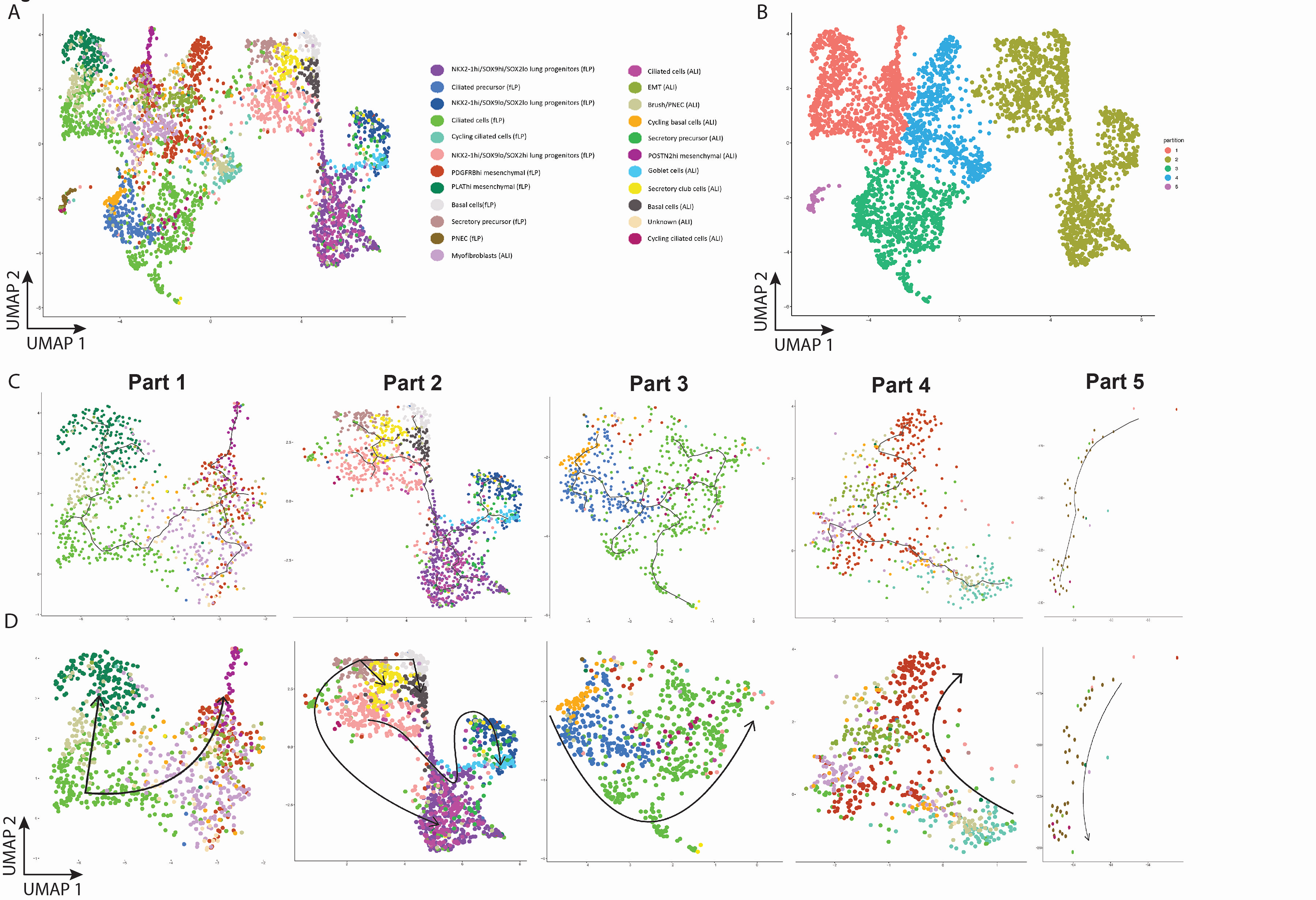
Pseudotime cell trajectory of differentiated cells from fetal to mature states. ***A.*** Integrated UMAP of fetal lung (fLP) and mature (ALI) cells calculated based on common anchor genes. ***B.*** UMAP visualization of partitioned clusters to determine lineage trajectories of cells. ***C.*** UMAP visualization of the pseudotime trajectories in each partition generated using *Monocle3*. **D.** RNA velocity trajectory, generated with *scVelo*, show changes in spliced RNA abundance of a given gene in a cell and determines the state of a cell based on these abundances over pseudotime. RNA velocity of the cells in each partition shows an asynchronous differentiation pattern of each cell with multiple originating nodes.

## MATERIALS & METHODS

### Embryonic stem cell (ES) and induced pluripotent stem cell (iPSC) expansion and maintenance

All human stem cell lines were approved for use by the Canadian Stem Cell Oversight Committee SCOC and the SickKids Research Ethics Board (REB # 1000071246 for use of human ES cells). Human ES line CA1 was provided courtesy of Dr. Andras Nagy (Lunenfeld Tanenbaum Research Institute, Toronto) and iPSC lines BU3-NKX2-1^GFP^ and BU3-NKX2-1^GFP^/P63^tdTomato^ (NGPT) were provided by Dr. Darrell Kotton (CReM Boston University, Boston). The hPSC lines were cultured in serum-free defined conditioned media mTeSR or mTeSR plus (STEMCELL Technologies) on hES-certified matrigel (Corning cat#354277). Briefly, cell culture plates were pre-treated with diluted Matrigrel (1:100 dilution) for at least 1 hour at room temperature or overnight at 4°C. Cells were grown to ∼80% confluency before passaging using ReLeSR^TM^ (STEMCELL Technologies, catalog #05872). Cells were split 1:6 with each passage. Media change was performed daily using mTeSR. Frozen vials of iPSC were thawed into mTeSR plus media supplemented with 10 µM ROCK inhibitor (Y27632, STEMCELL Technologies, catalog #72304) and the media changed after 2 days.

### Modified differentiation of iPSC into lung models

The new differentiation method was adapted and modified from our previously published protocol^18, 22^. Briefly, iPSCs are differentiated into definitive endoderm (Stage 1) using STEMCELL Technologies StemDiff DE kit (*Stage 1 media*), as per manufacturer’s protocol (catalog #05110). Definitive endoderm cells are then differentiated into anterior ventral foregut endoderm (Stage 2) using dual SMAD inhibition for 2 days with 2 µM Dorsomorphin (STEMCELL Technologies, cat#72102) and 10 µM SB431542 (STEMCELL Technologies, cat#72234) in media containing 500 ng/mL fibroblast growth factor-2 (FGF2, Preprotech, cat#100- 18B) and 50 ng/mL recombinant sonic hedgehog (SHH, Preprotech, cat#100-45) in Knockout DMEM (Gibco cat #10829-018), 10% Knockout serum replacement (KOSR, Gibco cat# 10828- 028), 1% penicillin/streptomycin, 2 mM Glutamax (Gibco, cat# 35050-061), 0.15 mM 1-mono- thioglycerol (MTG, Sigma-Aldrich, cat# M6145), and 1 mM non-essential amino acid (Gibco cat# 11140-050) (*Stage 2 media*). Cells were then switched to media containing only FGF2 and SHH for the remaining 3 days. On day 10 of differentiation (*Stage 3a media*), lung epithelial progenitors were induced using 10 ng/mL FGF7 (KGF, Preprotech, cat#100-19), 50 ng/ml FGF10 (Preprotech, cat#100-26), 10 ng/mL bone morphometric protein-4 (BMP4, R&D Systems, cat#314-BP), 3 µM CHIR99021 (STEMCELL Technologies, cat#72054), 100 nM All-trans Retinoic acid (ATRA; Sigma Aldrich cat# R2625), and 10 µM Y27632 for the first 2 days and without Rock inhibitor thereafter in lung basal media composed of IMDM/F12 (75:25), 0.5% N2 supplement-A (STEMCELLTechnologies, cat#07152), 1% B27 supplement (Thermo Fisher Scientific, cat#17504044), 2 mM Glutamax, 0.15 mM MTG, 0.05% bovine serum albumin (BSA, Thermo Fisher Scientific, cat#12483020), 50 µg/mL ascorbic acid (Sigma-Aldrich, cat# A4544), and 1% penicillin/streptomycin. On day 16 (*Stage 3b media*), proximalization of the lung progenitor cells was induced with media containing 50 ng/mL FGF7, 50 ng/mL FGF10, and 10 ng/mL FGF18 (Sigma-Aldrich, cat# F7301) in lung basal media for 5 days. On day 21 (*Stage 4 media*), media was switched to PneumaCult-Ex plus containing 1 µM forskolin, 10 ng/mL FGF18, 50 ng/mL FGF7, and 50 ng/mL FGF10 for an additional 5 days minimum. Cells can be cultured in this media for at least 10 days. To improve survival from the media switch, 10 µM Y27632 was added to the media for the first 2 days. On day 26 (*Stage 5a media*), stage 4 cells or airway organoids (AO) were seeded onto collagen-IV (Sigma Aldrich, catalog #C7521)-coated transwells (Thermo Fisher, Corning cat#01-200-161) at a seeding density of 2.5x10^5^ cells/insert. Cells were expanded to form a monolayer of epithelial cells on the transwell membrane prior to differentiation in Stage 5a media containing 2 µM Dorsomorphin, 10 µM SB431542, and 10 µM DAPT (Abcam, cat# ab120633) in PneumaCult-ALI media (without hydrocortisone, and supplemented with 50 nM dexamethasone, and 10 µM Y27632 for the first 2 days of culture). Cells were maintained in Stage 5a media on the apical side of the transwell until a monolayer of cells was formed. Once a confluent monolayer formed, media (*Stage 5b media)* comprised of PneumaCult-ALI media with hydrocortisone replaced with 50 nM dexamethasone, and supplemented 10 µM DAPT was added only to the basolateral side of the transwells to induce differentiation and polarization of the epithelium for up to 4 additional weeks.

Fetal lung cells at the end of stage 3 of the differentiation were sorted for pan-epithelial cell surface protein CD326 using magnetic activated cell sorting (MACS, Miltenyi) using LS columns (cat# 130-042-401). Non-epithelial (CD326-) cells were discarded and CD326+ single cells were seeded onto collagen IV-coated 96 well black/clear bottom plates (Thermo Fisher, cat# 165305) at a density of 25,000-50,000 cells/well. The following day, non-adherent cells suspended in media were removed and replaced with fresh Stage 3b media. Once the cells reached confluency (90%), CFTR functional studies via FLiPR were performed. CF iPSC-derived cultures were pre-treated with CFTR modulators prior to functional measurements.

To generate airway spheroids (AS), epithelial cells at the end of Stage 3 or Stage 4 were isolated by MACS as above. Single cells (CD326^+^) were resuspended in growth factor-reduced matrigel (BD354230) and immediately transferred to cell culture plates to form matrigel “domes” of 50 µL volume. Matrigel was allowed to solidify at 37 °C for 20 minutes before media was added, covering the dome for culture. For AS expansion of cells isolated from Stage 3, *Expansion Media* comprised of Pneumacult Ex Plus (STEMCELL Technologies, cat# 05040) supplemented with 1 µM A 83-01 (STEMCELL Technologies, cat# 72022) and 1 µM DMH-1 (STEMCELL Technologies, cat# 73632)^13^. For AS differentiation, *Expansion Media* was replaced with *Differentiation Media* comprised of Pneumacult-ALI complete media, 10 µM Y27632, 10 µM SB431542, 3 µM CHIR99021, 10 µM DAPT, 10 ng/mL FGF18, 10 ng/mL FGF7, 10 ng/mL FGF10, 1 µM forskolin, 1 µM IBMX, and 1 µM ATRA (Sigma Aldrich cat # R2625) for at least 7 days. Media change occurred every other day. Within 7 days, AS can be observed with characteristic bilayer concentric rings as observed in organoids derived from human primary bronchial epithelial cells. For swelling assays, 5000 CD326+ cells were seeded in semi-solid matrigel (50% matrigel in *Expansion Media*) on the apical side of polyester transwells. The mixture was allowed to solidify at 37 °C for 45 minutes before the bottom chamber was filled with 1.5mL of organoid media. Media change occurred every other day.

The above protocol was compared to directed differentiation protocols from our previous method^22^. Key differentiation stages were directly compared using two human PSC lines (human ES line CA1 and iPSC line BU3-NKX2-1^GFP^). At least 2 differentiation sets for each line were analyzed unless otherwise specified.

### Quantitative Real-time PCR

RNA was extracted using the RNeasy Kit (Qiagen, cat#74106) as per manufacturer’s protocol. Prior to RNA isolation, genomic DNA was removed using the QIAshredder (Qiagen, cat# 79656). Isolated RNA was reverse transcribed with Superscript II (Thermo Fisher Scientific, cat#18-064-071) as per manufacturer’s protocol. Quantitative real-time PCR was performed with PowerUP SYBR Green Mastermix Master Mix on ViiA7 (Applied Biosystems). Gene expression was normalized to house-keeping gene GAPDH and expressed relative to control human tissue (**Supplementary Table 1**) RNA extracts (2^ΔΔCT). Cycle threshold (CT) values above 38 were considered “not expressed”. Primer sequences are listed in **Supplementary Table 2**.

### In vitro Single cell RNA-sequencing (scRNAseq) preparation and sequencing

Samples were dissociated into single cell suspension with viability over 80%. For lung progenitor cultures, cells were washed twice with PBS, and dissociated with trypsin (Wisent, cat# 325-043- CL) at 37 °C for 4 minutes, then inactivated by a sorting solution, PBS containing 1% FBS. For ALI cultures, media was removed from both apical and basal chambers, and washed twice with PBS. Trypsin was added in both the apical and basal chambers, and incubated at 37 °C for 5 minutes. Trypsin in the basal chamber were removed, and cell pellets were collected with a sorting solution. For airway organoids, Matrigel was degraded by dispase (STEMCELL Technologies, cat# 07923) at 37 °C for 1 hour. After 2 PBS washes, trypsin was added to dissociate organoids at 37 °C for 5-10 minutes, then inactivated by sorting solution. After centrifugation at 300g for 5 minutes, the cell pellets were resuspended in sorting solution, and strained through 0.4 µm strainers (Falcon, cat# 352340). MULTI-seq barcoding was performed as per the protocol provided by Dr. Zev J Gartner Lab^52^. Cell number and viability were assessed by trypan blue (Gibco; cat# 15250061) with Countess II (Life Technologies, cat#A27977). Four Multi-seq barcoded samples were combined as one sample, and loaded 25,000 cells per GEM beads. GEM generation & barcoding, post GEM-RT cleanup & cDNA amplification, and Library construction steps were performed as per Chromium Next GEM single Cell 3’ Reagent Kits v3.1 (10x Genomics, cat#1000121, 1000120, 1000123). Sequencing was performed on the NovaSeq6000 at a read depth of approximately 60,000 reads/cell (TCAG Facility, SickKids).

### Preparation of single cell suspension from primary fetal lung tissue

GW 16 and GW18 fetal lung samples were procured from the Research Centre for Women’s and Infants’ Health (RCWIH) Biobank (SickKids REB# 1000067499 and LTRI REB #20-0035-E) and placed immediately in cold 1X PBS. Single cell suspension was achieved using the 37_C Multi_tissue_dissociation B program on the gentleMACS octo-dissociator (Miltenyi Biotec). Subsequently, cells were gently strained with 0.4 µm strainers. Lastly, cells were resuspended in RBC lysis buffer (Invitrogen, cat# 00-4333-57) for two minutes and cell viability was determined as described above. Single cell suspension was loaded at 6000 cells per GEM bead. GEM generation & barcoding, post GEM-RT cleanup & cDNA amplification, and Library construction steps were performed as per Chromium Next GEM single Cell 3’ Reagent Kits v3.1 (10x Genomics, cat#1000121, 1000120, 1000123). Sequencing was performed on the NovaSeq6000 at a read depth of approximately 60,000 reads/cell. (TCAG Facility, SickKids).

### Overview of single cell RNA-sequencing analysis pipeline

Single cell sequencing analyses were mainly performed with *Seurat v.4.0* as previously described^49^. Briefly, untrimmed reads were generated using *supernova/cellranger* mkfastq and bcl2fastq2 v2.20. Reads alignment to human reference genome and generation of gene expression matrix were done using 10x Genomics *Cell Ranger v6.0.1* software with provided hg19 (GRCh38) reference genome and transcriptome annotation. MULTI-seq barcodes were demultiplexed using *deMULTIplex*^52^, an available R package provided by the Gartner Lab. In each sample, to ensure high data quality for further analysis, cells with more than 20,000 or less than ∼1,000 genes, or mitochondrial transcript fraction over 10%, were filtered out. Genes expressed in less than 3 cells were excluded from our analysis. Cell cycle phase was scored by expression levels of cell cycle related genes. Cellular variance of total number of UMI, mitochondrial transcript fraction, and cell cycle phase was regressed out and gene expression levels were normalized via *SCTransform*^53^. For the *in-vitro* cultures, similar lung differentiation cultures derived from different cell lines were merged into one dataset through the Seurat’s *merge* function. In the merged samples, principal component analysis (PCA) was performed based on normalized (Pearson residuals) expression levels of genes. Graph-based (Louvain) clustering approach and UMAP dimension reduction was performed based on between the top 50 principle components. Primary fetal lung datasets were subsetted for cells expressing EPCAM for correlation analysis. Identification of positive markers for each cluster was performed with the two-sided Wilcoxon rank sum test^54^. Differential gene expression analysis was used to manually annotate based on previously published work^5–11, 13, 36, 55^. Each *in-vitro* differentiation culture was analyzed separately. To quantify the transcriptome similarity between *in-vitro* cells and control fetal lung cell sub-clusters, Pearson’s correlation coefficients of expression levels were calculated using shared genes of primary and *in-vitro* cells. Gene ontology enrichment in cluster markers was performed using *DAVID v6.8*^56, 57^. Significant enrichment was defined as Benjamini-corrected P < 0.05. Pseudotime analysis of samples were performed using *Monocle3*^48^. In brief, datasets from fetal lung and air-liquid interface cultures were integrated using *Seurat*’s integration pipeline^47^. The resulting dataset was divided into partitions^50^ and its trajectories were subsequently analyzed through Monocle3’s *learngraph()* function. RNA velocity analyses (performed using sc*Velo*) were estimated based on the rate of gene expression change for an individual gene at a given time based on the ratio of its spliced and unspliced messenger RNA (mRNA)^51^. In particular, the stochastic model assumes transcription and splicing events as probabilistic events using second-order moments (representing the variation of unspliced and spliced mRNA), rather than deterministic ordinary differential equation model that does not take into account gene expression stochasticity. RNA velocities were computed for every partition generated by Monocle3. Velocities were projected using scVelo’s scv.pl.velocity_embedding_stream and scv.pl.velocity_embedding functions. Bioinformatics code is available on our github: https://github.com/SickKids-Wong-Lab/Ngan-2021--Protocol-Comparison-Paper.

### Flow cytometry

Samples were dissociated with TrypLE for 5 minutes at 37 °C, and collected at 300g for 5 minutes. The cells were washed once with 3 ml of FACS buffer (PBS -Ca^2+^/Mg^2+^ with 0.5% BSA), and then transferred to a FACS tube containing a filter top to remove cell clumps. Single cell suspensions in 200 ul FACS buffer were stained with Fixable Viability Dye eFluor450 (eBioscience, cat# 65-0863-14) for 30 minutes on ice. Cells were then washed 3 times prior to fixation with 4% PFA for 10 minutes at room temperature. Primary antibodies (**Supplementary Table 3**) were added to resuspended fixed cells in 200ul FACS staining buffer at 4 °C for 30 min followed by secondary antibody staining (if needed) at 4 °C for another 30 mins. After a brief wash, the cells were resuspended in FACS buffer and analyzed with the BD Biosciences LSRII analyzer. Data analysis was performed using the FlowJo software.

### Immunofluorescence

Monolayer cultures were washed twice with PBS (without Ca^2+^ and Mg^2+^) and then fixed with ice-cold methanol for 20 minutes, followed by another round of wash with PBS. Cells were subsequently permeabilized with 0.25% Triton X-100 for 20 minutes. The cells were then treated with a blocking solution containing 5% BSA and 0.25% Triton X-100 to block non-specific binding at room temperature for 30 minutes to 1 hour. Primary antibodies (**Supplementary Table 3**) were added to the blocking solution and incubated overnight at 4℃. Cells were then washed 2X with PBS followed by incubation with secondary antibodies (**Supplementary Table 3**) at room temperature for 2 hours. Finally, another wash was performed and the nuclei counterstained with DAPI for 15 minutes at room temperature. Stained cells were kept in PBS and images captured using the Olympus Spinning Disc Confocal within 2 days of staining.

Organoids were released from the matrigel bubble using ice cold dispase (STEMCELL Technologies, cat# 07923) and kept on ice for about 1 hour with periodic mixing to gently melt the matrigel. The organoid suspension was then centrifuged at 300 g for 3 minutes to pellet the organoids. After careful removal of the melted matrigel in the dissociation buffer, the organoids were resuspended in fixative (4% PFA) for 15 minutes at room temperature. After a brief wash with PBS containing 0.25% TritonX-100, the organoids were resuspended in PBS containing 5% BSA and 0.1% TritonX-100 and kept at 4 ℃ until staining. For organoid staining, organoids were incubated with primary antibodies for overnight incubation at 4 ℃. After two washes with PBS, secondary antibodies and Hoescht dye (1:1000, Thermo Fisher Scientific, cat# H3570) were added for 1-2 hour incubation at room temperature. The organoids were washed with PBS twice before mounting on superfrost slides (Fisher Scientific, cat# 22-037-246). 30ul droplets containing the organoids were gently placed onto the slide and pre-mounted with 0.12mm Secure-Seal silicone spacers (Thermo Fisher cat# S24737) before securing into place with a coverslip. Organoids were imaged within 2 days of staining and stored in a humidified chamber at 4 ℃ protected from light leading up to it. Fluorescence images were captured with the Olympus spinning disc confocal microscope and analyzed with the Volocity imaging software (Quorum Technologies).

For fixed paraffin-embedded slide staining of ALI cultures, slides were first deparaffinized and antigen retrieval was performed using either citrate buffer or HIER solution (Abcam, ab208572). Successive steps were followed as previously described^22^. Primary adult lung slides (Amsbio, cat# T2234152) were used as staining controls.

Isotype controls using non-immune immunoglobulins were used to assess non-specific binding and background fluorescence for imaging.

### Air-liquid Interface Apical chloride conductance (ACC) Assay for CFTR function

The ACC assay was used to assess CFTR mediated changes in membrane depolarization using methods as previously described^23^. For ACC measurements, basal chamber were filled with HBSS (Wisent, cat# 311-513), and the cells on transwells were incubated with zero sodium, chloride and bicarbonate buffer (NMDG 150 mM, Gluconic acid lactone 150 mM, Potassium Gluconate 3 mM, Hepes 10 mM, pH 7.42, 300 mOsm) containing 0.5 mg/ml of FLIPR® dye for 30 mins at 37 °C. CFTR function was measured after acute addition of Fsk (10 µM) or 0.01% DMSO control. CFTR functional recordings were measured using the SpectraMax i3. Modulators were then added to stimulate CFTR mediated anion efflux and CFTR mediated fluorescence changes were monitored and imaged every 2 minutes. CFTR channel activity was terminated with addition of CFTR-inh172 (10 µM) and fluorescence changes were monitored every 2 minutes for 5 reads.

### Submerge culture apical chloride conductance assay

The ACC assay was used to assess CFTR mediated changes in membrane depolarization using methods as previously described^28^. iPSC-derived airway cultures were replated on collagen IV (Sigma Aldrich, cat# C7521)-coated 96 well plates (Corning, cat# 3603) at AFVE stage Day 3. Cells were dissociated using TrypLE for 3 minutes at 37 °C, and replated at 25,000 cells/well in AFVE media. The AFVE cells continued differentiation until S3B stage. For ACC measurement, the cells on 96 well plates were incubated with zero sodium, chloride and bicarbonate buffer (NMDG 150 mM, Gluconic acid lactone 150 mM, Potassium Gluconate 3 mM, Hepes 10 mM, pH 7.42, 300 mOsm) containing 0.5 mg/ml of FLIPR® dye for 30 mins at 37 °C. CFTR function were measured after acute addition of FSK (10 µM) or 0.01% DMSO control. CFTR functional recordings were measured using the FLIPR® Tetra High Throughput Cellular Screening System (Molecular Devices), which allowed for simultaneous image acquisition of the entire 96 well plate. Images were first collected to establish baseline readings over 5 mins at 30 second intervals. Modulators were then added to stimulate CFTR mediated anion efflux and CFTR mediated fluorescence changes were monitored and images were collected at 15 second intervals for 70 frames. CFTR channel activity was terminated with addition of CFTR-Inh172 (10 µM) and fluorescence changes were monitored at 30 second intervals for 25 frames.

### Split-open apical chloride conductance (ACC) Assay for CFTR function

The ACC assay was used to assess CFTR mediated changes in membrane depolarization using methods as previously described^40^. Briefly, AO were “split open” by plating them on 96 well assay plate (Corning; cat# 3603) and allowed to adhere and “hatch” for up to 3 days. At this point, a layer of epithelia can be observed adhered to the transwell membrane. For ACC measurements, the cells on transwells were incubated with zero sodium, chloride and bicarbonate buffer (NMDG 150 mM, Gluconic acid lactone 150 mM, Potassium Gluconate 3 mM, Hepes 10 mM, pH 7.42, 300 mOsm) containing 0.5 mg/ml of FLIPR® dye for 30 mins at 37 °C. CFTR function were measured after acute addition of FSK (10 µM) or 0.01% DMSO control. CFTR functional recordings were measured using the FLIPR® Tetra High Throughput Cellular Screening System (Molecular Devices), which allowed for simultaneous image acquisition of the entire 96 well plate. Images were first collected to establish baseline readings over 5 mins at 30 second intervals. Modulators were then added to stimulate CFTR mediated anion efflux and CFTR mediated fluorescence changes were monitored and images were collected at 15 second intervals for 70 frames. CFTR channel activity was terminated with addition of CFTR-inh172 (10 µM) and fluorescence changes were monitored at 30 second intervals for 25 frames.

### Spheroid swelling assay

Spheroids were cultured in semi-solid Matrigel (50% matrigel in *Expansion Media*)in transwells for up to 10 days, until a lumen was observed. On the day of swelling, organoids were released from the matrigel using dispase on ice for 1-2 hours with intermittent mixing in between. Floating spheroids were then transferred to an ultra-low adhesion cell culture plate to culture in suspension two days prior to FIS assay in forskolin-free airway organoid media. For FIS, suspended spheroids were transferred to fresh media containing 10 µM of forskolin and 1 µM IBMX and imaged immediately to capture T0 size. At 4-6 and 24 hours post stimulation, images of the spheroids were also captured for size measurements. The diameter of organoids were calculated using the imageJ software.

### Immunoblotting

Cells and organoids were lysed in modified radioimmunoprecipitation assay (RIPA) buffer (50 mM Tris-HCl, 150 mM NaCl, 1 mM EDTA, pH 7.4, 0.2% SDS (Invitrogen, cat# 15553), and 0.1% Triton X-100 (Sigma Aldrich, cat#93443) containing a protease inhibitor cocktail (GBiosciences, cat# 031M-D)^58^. Soluble fractions were run by SDS-PAGE on 6% tris-glycine gels (Life Technologies, cat# XP00065BOX). Electrophoresis were run in Tris-glycine-SDS buffer (Wisent, cat# 811570) at 100V for up to one and a half hours. Proteins were transferred to nitrocellulose membranes (Bio-Rad) in Tris-glycine buffer (Wisent, cat# 811560) with 20% methanol (Fisher Scientific, cat# 67-56-1) at 100V for one hour, then blocked with 5% milk solution for half an hour at room temperature. Transferred blots were stained with primary antibodies (**Supplementary Table 2**) overnight at 4 °C, followed with secondary antibodies for 1 hour at room temperature. Signal was developed using Clarity Max Western ECL Substrate (Bio-Rad, cat# 1705062) and captured by Licor Odyssey Fc Imaging System. CFTR protein level was normalized to calnexin protein level analyzed by ImageStudio Lite (Li-COR Biosciences).

## ACKNOWLEDGEMENT

We would like to thank Drs. D. Kotton and F. Hawkins (Boston University) for providing the human iPSC reporter lines and Dr. Nagy (Lunenfeld Tanenbaum Research Institute, Toronto) for the human ESC CA1 line. Monoclonal CFTR antibodies #562 were courtesy of J.R. Riordan (University of North Carolina, Chapel Hill, North Carolina, USA). Small molecule CFTRinhibitor-172 was obtained from CFFT (Cystic Fibrosis Foundation Therapeutics). This work was supported by the CF Canada-SickKids Program in Individual Therapy (CFIT) (in part) by SickKids CFIT Microsynergy grant (awarded to A.P.W in 2017) and Vertex Pharmaceutical CF Research Innovation Award (awarded to A.P.W in 2017), the Genome Canada and the Ontario Genomics Institute (OGI-148), the SickKids Foundation & CIHR-IHDCHY grant (NI20-1070), and the Stem Cell Network Early Career Investigator - Innovation Award (FY21/ECI23 Wong). Computational power was enabled by support provided by Compute Canada (www.computecanada.ca) for processing scRNA-seq datasets. We would like to thank Drs. Chris McGinnis and Zev J Gartner (University of California San Francisco, U.S.A) Lab for providing the reagents and protocols for MULTI-seq experiment.

## AUTHOR CONTRIBUTIONS

S.Y.N, H.Q, J.D, J.L, O.L, S.X, E.H, A.P.W performed experimental assays, analyzed the data, and contributed to the preparation of the manuscript. S.Y.N, and H.Q contributed equally to the work. A.P.W, S.Y.N, and H.Q designed the study, interpreted the data and wrote the manuscript. All authors read and approved the manuscript.

## DISCUSSION

Human PSCs have been an invaluable resource in generating human lung tissues for regenerative medicine and development. While there are many protocols available to date that use common factors to direct the differentiation of hPSC to lung epithelial cells, incremental improvements have led to the ability of generating distal versus proximal lung epithelial cell types, mature cell types, and renewable progenitors^13–15^. The latter is especially important for generating a renewable source of lung epithelial cells from iPSC without having to repeat a lengthy and costly differentiation protocol from PSC state. We previously were the first to report a method to generate airway epithelial cells from human pluripotent stem cells that can be used to model cystic fibrosis disease *in-vitro*^18^. However, without fractionation or enrichment for putative lung progenitor cells during the early stages of differentiation, non-lung epithelial lineages co-emerged throughout the differentiation process, not surprisingly since many of the morphogenetic pathways are shared during development. Few groups have shown efficient generation of NKX2-1^+^ lung progenitor (>90%)^26, 30^, while others have relied on using reporter strategies (NKX2-1^GFP+^, NKX2- 1^GFP+^SFTPC^tdTomato+^, NKX2-1^GFP+^P63^tdTomato+13–15or^ identified putative cell surface markers (CD47^high^ and CD26^low^ or CPM^+^)^14, 24, 25, 39^ to isolate or enrich for the lung progenitor population. Nonetheless, significant improvements in purifying lung epithelial progenitor cells have provided an enriched source of cells to model lung diseases *in-vitro*^11, 13, 14^. Temporal activation of Wnt pathway is key in directing differentiation of distal type II alveolar epithelial cell (AEC2)^14, 15^ and can be used to model distal respiratory diseases such as surfactant deficiencies. Inhibition of the pathway promotes proximal cell differentiation and can be used to generate proximal epithelial organoids to model CF airway disease *in-vitro*^14^.

Recently, Hawkins et al^13^ showed a robust method of generating renewable sources of basal stem cells with the capacity to differentiate into multi-epithelial cell types, including secretory and ciliated cells *in-vitro,* and repopulate the denuded rat trachea *in-vivo*. Using single cell analyses of primary fetal lung tissue of the pseudoglandular stage, Miller et al identified SMAD signalling as a key regulator of distal tip-to-airway transition of the epithelial cells, and in manipulating this signalling, will generate renewable fetal basal stem cells with multilineage differentiation potential^11^. Here, we present a similar approach of generating a renewable progenitor population enriched in basal stem cells that improves the differentiation efficiency in generating fetal-like lung cells and mature airway epithelia in ALI. Similar to previous protocols, dual SMAD inhibition using small molecules A83-01 and DMH-1^13^ in addition to CHIR99021 (Wnt pathway activation) and all-trans retinoic acid (RA pathway activation)^25, 30^ generates epithelial organoids with expansion potential. Employing single cell RNA sequencing to benchmark and determine the heterogeneity of the cell types in our differentiation cultures, we identified unique clusters of lung cells found in the developing fetal lung^31^, as well as the co-emergence of mesenchymal cells. The origin of the mesenchymal cell population remains unclear, though we speculate that epithelial-mesenchymal transition may contribute to these cell populations^45^. Indeed, gene ontology assessment of differentially expressed genes (DEG) of each cell cluster revealed high activity of epithelial to mesenchymal transition (EMT and myofibroblast) and positive regulation of epithelial to mesenchymal transition (myofibroblast and POSTN2^hi^ mesenchymal) in the top GO terms.

Limited access to primary human fetal tissues for biomedical research has made it particularly difficult to study cellular and molecular mechanisms driving human lung development. Much of what we know about lung development has traditionally been extrapolated from heralding efforts in animal models^4, 59, 60^ and only recently been studied in limited human tissue samples^31, 61^. These models have been useful in describing conserved developmental pathways. However, not all pathways are conserved and human disease development is not always recapitulated in animal models, as in the case of CF^60^. Human stem cell-derived models have gained increasing traction to study human-specific developmental pathways and disease modeling^13, 26, 62–65^. As the differentiation of hPSC follows a directed developmental trajectory, these new human tissue models are of great interest to study fundamental mechanisms driving lung cell lineage formation. However, many of the hPSC-derived “recapitulated’’ developmental stages have largely been validated with few gene expression markers benchmarked from previous mouse studies including our previous developmental protocol^18, 22^. Few studies have employed human primary tissue to validate stage-specific match of the cells formed from *in-vitro* differentiation using RNA sequencing (single cell or bulk) of *select* primary tissues^11, 15^. Current challenges of *in-vitro* human pluripotent stem cell (hPSC)-derived differentiations towards tissue-specific cell types involve the faithful modeling of *in-vivo* development. In our study, we generate fLP reminiscent of GW16 pseudoglandular fetal lung that contain cells with unique SOX2 and SOX9 expression patterns similar to human fetal bud tip proximal-distal patterning^31^. This provides a unique hPSC-derived lung model to assess cell patterning and specification during fetal lung development and can potentially be used to determine genetic causes of congenital pulmonary airway malformations^66^. This is particularly important as access to human fetal tissues for research is limited and exacerbated by the scarcity of fetal lungs with congenital abnormalities. We confirmed high expression of CFTR, an anion channel protein implicated in CF disease, in our fLP cultures which has previously been found to be up-regulated during pseudoglandular lung development^32^. We demonstrated measurable CFTR function in fLP cultures, which suggests a developmental role of CFTR in fetal lung cells that may currently be underappreciated. Thus, we have now identified a potential opportunity to understand the congenital cause of CF lung diseases using hPSC-derived fLP models, as previous studies have found pathological manifestations of CF diseases in newborns^67–72^.

A new advantage of our improved differentiation protocol is the generation of expandable hPSC-derived airway spheroids (AS) from fLEC. Similar to previous findings^11, 13^, dual SMAD inhibition appears to be enriched for basal stem cell-like cells expressing TP63 that are capable of multilineage differentiation *in-vitro*. Importantly, the hPSC-derived AS model provides a user-friendly opportunity that is time and cost-effective for researchers with no experience or capacity to work with hPSC. Their proliferative and relatively undifferentiated nature enables rapid expansion of AS for regenerative studies^13^ and developing high throughput assays for disease models and/or drug screens^28^. Upon differentiation, the airway epithelial organoids recapitulate proximal airway phenotype with basal cells on the basolateral side and secretory and ciliated cell types localized to the luminal side. CFTR is expressed on the luminal membrane of these cells. While some studies have suggested airway organoids can be used in forskolin-induced swelling (FIS) assays to assess CFTR function^14^, the slow (>4 hours of FIS) and typically minimal swelling (often <10%) observed in control AS, likely due to much lower expression of CFTR, limits the use of AS in rapid screens FIS screens of CFTR modulators, unlike intestinal organoids in which CFTR is more abundantly expressed^29, 73^. Instead, our open AS apical chloride conductance (ACC) assay is much more sensitive in detecting CFTR-mediated responses and offers an alternative strategy to measure CFTR modulator responses in CF iPSC-derived AS. It should be noted that an additional limitation of AS is the relatively simple epithelium that is formed devoid of other airway epithelial cells types, including rare cell types such as neuroendocrine cells and ionocytes.

Therefore, contributions of the rare ionocytes in CF disease and responses to therapies are not captured with the AS model.

Similar to Hawkins et al^13^, further differentiating the cells in ALI as we previously described^18, 22^ induces maturation and polarization of the epithelium, such that by the end of 5 weeks in ALI, a pseudostratified epithelium is formed, reminiscent of the large airways. Our hPSC-derived ALI cultures showed improved levels of CFTR protein and measurable function (up to 40% function above baseline). Most importantly, air-liquid interface mimics post-natal airway development. In our study, we used scRNAseq to reveal transitory cell types that could not be captured previously^22^. Many of the known airway cell types, including ciliated, basal, goblet, club, PNEC, and ionocytes, were identified in our ALI cultures, in addition to a large population of mesenchymal cells that co-developed during cellular differentiation. Interestingly, an unknown cluster of cells appeared to emerge from cells undergoing EMT and express some genes (*IQGAP2, ID3, ID4*) previously associated with ionocytes but not *FOXI1*^8^. Moreover, CFTR is not highly expressed in this cluster of cells, suggesting these may be pre-ionocyte cell types and that our current differentiation protocol does not support ionocyte cell development.

Previously, pulmonary cell types were identified through a snapshot measuring the most abundant genes at each stage of the differentiation protocol^18, 22^. In our current integrated analyses of the fLP and ALI differentiation stages through functional pseudotime analysis, a trajectory inference method that models gene expression changes as a function of pseudotime along lineages^48, 74^, we were able to identify lung cell lineage hierarchies and relationships as they differentiate in culture. We found previously identified lineage relationships, such as basal cells as the source of secretory and ciliated cells^75^. We also identified the emergence of the embryonic bud (NKX2-1^hi^SOX9^hi^SOX2^lo^) that gives rise to stalk (NKX2-1^hi^SOX9^lo^SOX2^hi^) progenitor cell types and the presence of NKX2-1^hi^SOX9^lo^SOX2^lo^ cells reminiscent of the developing saccule as previously found^31, 75^. Interestingly, we also identified ciliated cells as a source of mesenchymal cells, which supports previous studies demonstrating a downregulation in CFTR and concomitant E-cadherin induced EMT phenotypic changes in cancer and CF^2,76^. Interestingly, we also found overlaps of cell clusters such as NKX2-1^hi^SOX9^hi^SOX2^lo^ lung progenitors with ALI ciliated cells and ALI cycling basal cells with ciliated precursors in the fetal stage, suggesting common transcriptional programs shared between mature and fetal lung stages. An isolated cluster of PNEC that emerged in the differentiation shared no common gene expression to other cell types and may have emerged from a separate ancestral lineage that has not been captured in our cultures^77^. Overall, we identified unique lineage relationships between the cells as they emerge in our differentiation between fetal and mature cell states.

We present a new and improved hPSC differentiation protocol that models human lung cell development *in-vitro* and may be used to study early developmental mechanisms driving congenital lung disease pathogenesis. Furthermore, creating new pre-clinical tools to investigate signalling mechanisms driving development, that can reactivate during airway regeneration, can provide insight into activating novel endogenous pathways for cell replacement therapy.

## Notes

### Competing Interest Statement

The authors have declared no competing interest.

### Summary of Updates

This version of the manuscript has been revised to clarify the phenotype of the cells generated in the differentiation especially at stage 3. In addition, grammatical and spelling errors have been fixed in this version.

