## Supplementary files for "Modeling lung cell development using human pluripotent stem cells"

**Supplementary Fig. 1.** Comparison of protein and mRNA expression levels of cultures at the anterior ventral foregut endoderm stage (Stage 2) for new and old differentiation protocols. ***A.*** Flow cytometric analysis for anterior foregut (SOX2), midgut (PDX1) and hindgut (CDX2) cells. Red contour plot is stained for respective lineage markers and the grey contour plot represents unstained control. ***B.*** Gene expression analysis by real-time quantitative reverse transcription polymerase chain reaction (qRT-PCR). Genes were normalized to housekeeping gene *GAPDH* and expressed relative to RNA from fetal lung (whole lung from gestational week 18 tissue). At least 3 technical replicates were performed, n= 2 biological (previously published human pluripotent stem cell lines: CA1 and BU3-NKX2-1$^{GFP}$). Bar represents the mean expression level.

**Supplementary Fig. 2.** Comparison of gene expression levels of cultures at fetal lung progenitor stage (fLP). Genes were normalized to housekeeping gene *GAPDH* and expressed relative to RNA from fetal lung (whole lung from gestational week 18 tissue). At least 3 technical (differentiation) replicates were performed, n= 2 biological (previously published human pluripotent stem cell lines:CA1 and BU3-NKX2-1$^{GFP}$). Bar represents the mean expression level.

**Supplementary Fig. 3.** Characterization of fetal lung progenitors (fLP) cells generated with our old differentiation protocol. ***A.*** Dot plot shows the average gene expression level and frequency of expressed genes in each cluster of fLP cells derived from the old protocol. ***B.*** Heatmap represents the top (up to) 5 gene ontology (GO) terms identified with each cluster based on differential gene expression (DEG). Empty boxes represent no or insignificant p-values. ***C.*** Immunohistochemistry staining for TTF1 (lung), SOX9 (distal), SOX2 (proximal), CFTR and beta-catenin show the presence of these markers in fLP cells derived from two independent hPSC lines (hES: CA1 and hiPSC: BU3-NKX2-1$^{GFP}$ lines).

**Supplementary Fig. 4.** Comparison of bulk gene expression levels of cultures after 5 weeks of ALI (Stage 5). Genes were normalized to housekeeping gene *GAPDH* and expressed relative to RNA from adult lung (normal 39yo lung tissue) or trachea tissue (normal 29yo). At least 3 technical (differentiation) replicates were performed, n= 2 biological (previously published human pluripotent stem cell lines: CA1 and BU3-NKX2-1$^{GFP}$). Bar represents the mean expression level.

**Supplementary Fig. 5.** Single cell RNA sequencing analysis of ALI cultures using the old differentiation protocol. ***A.*** Dot plot of the average gene expression level and frequency of expressed genes in each cluster of ALI cells. ***B.*** Heatmap represents the top (up to) 5 gene ontology (GO) terms identified with each cluster based on differential gene expression (DEG). The Benjamini-adjusted p-value in log scale (<0.05 or log (-1.3)) was used to determine DEG. Empty boxes represent no or insignificant p-values.

**Supplementary Table 1**: Positive control for quantitative PCR

| Tissue | Company | Catalog # | Lot # | Age | Gender |
|---|---|---|---|---|---|
| Adult Lung | Cell Applications Inc. | 1H-40 | 2788 | 39 | Male |
| Fetal Lung | Cell Applications Inc. | 1F-40 | 1395 | Gestational week 18 | NA |
| Trachea | BioChain | R1234160 | B83066 | 29 | Male |

**Supplementary Table 2**: List of Primers

| Gene | Forward (5'-->3') | Reverse (5'-->3') |
|---|---|---|
| ASCL1 | GGGCTCTTACGACCCGCTCA | AGGTTGTGCGATCACCCTGCTT |
| CCSP (SCGB1A1) | TTCAGCGTGTCATCGAAACCCC | ACAGTGAGCTTTGGGCTATTTTT |
| CFTR | CTATGACCCGGATAACAAGGAG G | CAAAAATGGCTGGGTGTAGGA |
| FOXA2 | GGGAGCGGTGAAGATGGA | TCATGTTGCTCACGGAGGAGTA |
| FOXG1 | CTCCGTCAACCTGCTCGCGG | CTGGCGCTCATGGACGTGCT |
| FOXI1 | CGGGCAAAGGGAATTACTGG | AGGCTCCATCCAAGATGTCC |
| FOXJ1 | GAGCGGCGCTTTCAAGAAG | GGCCTCGGTATTCA CCGTC |
| FOXP2 | AATCTGCGACAGAGACAATAAG C | TCCACTTGTTTGCTGCTGTAAA |
| GAPDH | ACAACTTTGGTATCGTGGAAGG | GCCATCACGCCACAGTTTC |
| KRT5 | GGAGTTGGACCAGTCAACATC | TGGAGTAGTAGCTTCCACTGC |
| KRT8 | GGTGGACCCCAACATCCA | GCTGCAGGAGGCTCCACTT |
| KRT13 | AGGACGCCAAGATGATTGGTT | GTGGTAACAGAGGTGCTACGG |
| KRT18 | CAGAGATCGAGGCTCTCAAGGA | GTCAACCCAGAGCTGGCAAT |
| MUC5ac | CCATTGCTATTATGCCCTGTGT | TGGTGGACGGACAGTCACT |
| SCGB3A2 | CGGAATTCCCCAGATAACTGTCA | ACATCTAGACACCAAGTGTGAT AGC |
| SFPTC | CACCTGAAACGCCTTCTTATCG | TGGCTCATGTGGAGACCCAT |
| SOX2 | TACAGCATGTCCTACTCGCAG | GAGGAAGAGGTAACCACAGGG |
| SOX9 | GAGGAAGTCGGTGAAGAACG | CCAACATCGAGACCTTCGAT |

| | | |
|---|---|---|
| TFF1 | CCCCGTGAAAGACAGAATTGT | GGTGTCGTCGAAACAGCAG |

**Supplementary Table 3**. List of Antibodies

| Primary Antibodies | Isotype | Host | Company | Catalog # or Clone ID | Dilution | Assay |
|---|---|---|---|---|---|---|
| Beta-tubulin | IgG | Rabbit | Abcam | Ab15568 | 1:200 | IF |
| Calnexin | IgG | Rabbit | Sigma Aldrich | C4731 | 1:2500 | WB |
| CFTR | IgG2b | Mouse | courtesy of J.R. Riordan | 596 | 1:1000 | WB |
| CFTR | IgG1 | Mouse | Millipore | 05-581 | 1:500 | IF |
| CGRP | IgG | Mouse | Sigma | C9487 | 1:200 | IF |
| deltaN P63 | IgG | Rabbit | BioLegend | 619002 | 1:200 | IF/FC |
| E-cadherin | IgG | Rat | BioLegend | 147302 | 1:500 | IF |
| FOXI1 | IgG | Goat | Sigma | SAB2501502 | 1:200 | IF |
| FOXJ1 | IgG | Mouse | Invitrogen | 149965-82 | 1:200 | IF/FC |
| KRT5 | IgG | Rabbit | Abcam | Ab53121 | 1:200 | IF |
| KRT14 | IgG3 | Mouse | Abcam | ab7800 | 1:200 | IF |
| KRT8/18 | IgG | Rabbit | Abcam | ab53280 | 1:200 | IF |
| MUC5ac | IgG | Mouse | Thermo Scientific | MA5-12178 | 1:200 | IF |
| pan-cytokeratin | IgG1 | Rabbit | Abcam | ab234297 | 1:500 | IF |
| PDX1 | IgG | Mouse | Santa Cruz | sc-390792 | 1:500 | FC |
| SCGB3A2 | IgG | Goat | Santa Cruz | sc-48320 | 1:200 | IF |
| SOX2 | IgG | Goat | R&D | AF2018 | 1:200 | IF/FC |
| SOX9 | IgG2a | Mouse | Abcam | ab79976 | 1:200 | IF |
| TTF1 | IgG | Rabbit | Abcam | ab76013 | 1:100 | IF/FC |

| | | | | | | |
|---|---|---|---|---|---|---|
| Alexa Fluor 647 Phalloidin | | | Invitrogen | A22287 | 1:500 | IF |
| ZO-1 | IgG1k | Mouse | Invitrogen | 33-9100 | 1:200 | IF |
| **Secondary Antibodies** | **Isotype** | **Host** | **Company** | **Catalog # or Clone ID** | **Dilution** | **Assay** |
| Anti-rabbit IgG (H+L) Alexa Fluor 488 | IgG | Donkey | Invitrogen | A11008 | 1:500 | IF |
| Anti-mouse IgG (H+L) Alexa Fluor 546 | IgG | Donkey | Invitrogen | A10036 | 1:500 | IF |
| Anti-goat IgG (H+L) Alexa Fluor 568 | IgG | Donkey | Invitrogen | A11057 | 1:500 | IF |
| Anti-mouse IgG (H+L) Alexa Fluor 647 | IgG | Donkey | Invitrogen | A31571 | 1:500 | IF |
| Anti-mouse IgG (H+L)-HRP conjugate | IgG | Goat | BIO-RAD | 170-6516 | 1:700 | WB |
| Anti-rabbit IgG (H+L)-peroxidase | IgG | Goat | Jackson ImmunoResearch | 111-035-003 | 1:5000 | WB |

**Supplementary Table 4: List of small molecules**

| Small molecule | Source | Catalog # | Final concentration |
|---|---|---|---|
| 3-Isobutyl-1-methylxanthine | Sigma Aldrich | I5879 | 1 μM |
| CFTR inhibitor-172 | Cystic Fibrosis Foundation | B1 | 10 μM |

| | | | |
|---|---|---|---|
| A83-01 | Sigma Aldrich | SML0788 | 1 μM |
| CHIR99021 | STEMCELL Tech | 72054 | 10 μM |
| DAPT | Abcam | ab126033 | 10 μM |
| DMH-1 | Tocris | 73632 | 1 μM |
| Dorsomorphin | STEMCELL Tech | 72102 | 2 μM |
| Forskolin | STEMCELL Tech | 72114 | 10 μM |
| SB431542 | STEMCELL Tech | 72234 | 10 μM |
| Y27632 | STEMCELL Tech | 72304 | 10 μM |

**Supplementary Fig. 1**

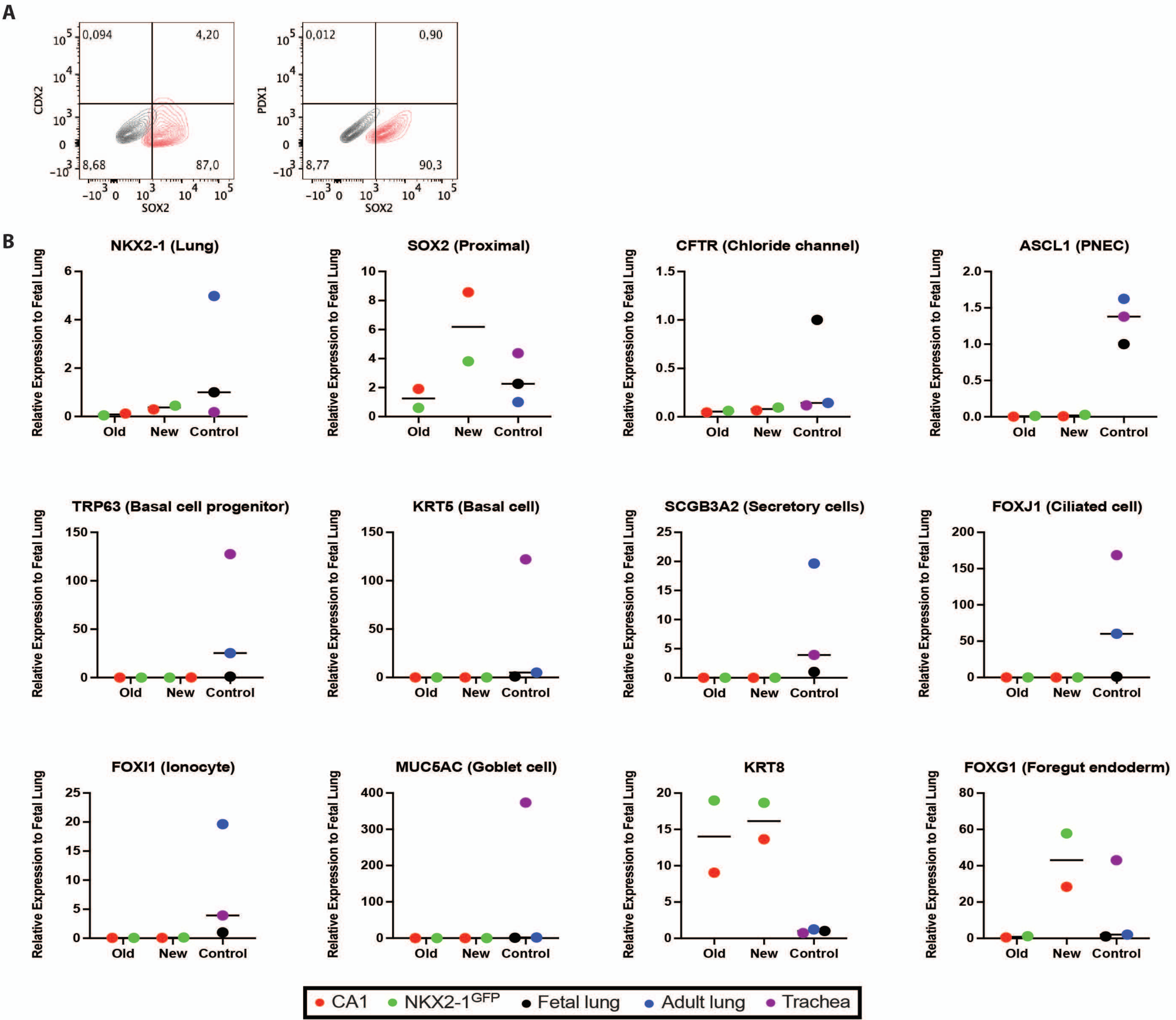

Supplementary Fig. 2

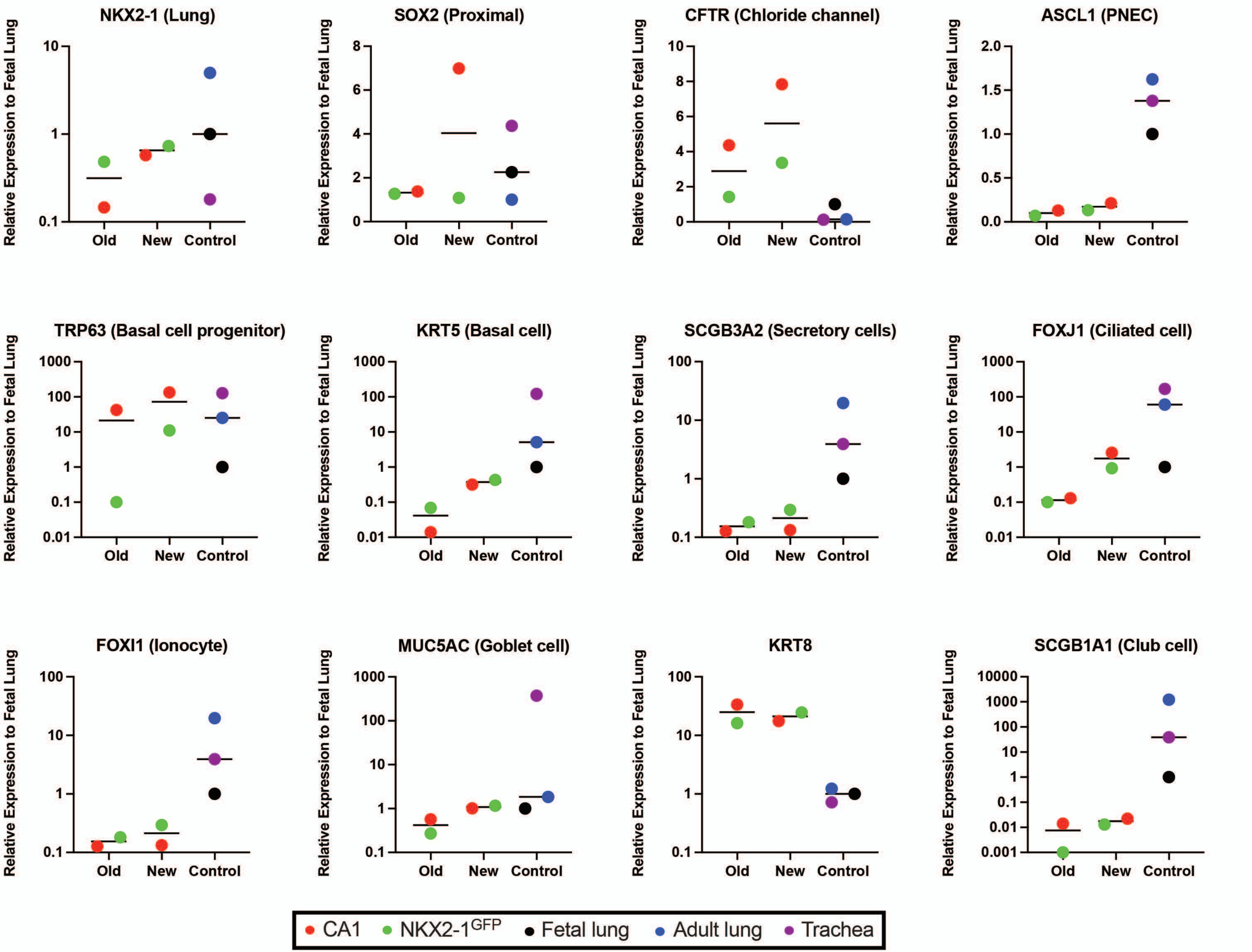

Supplementary Fig. 3

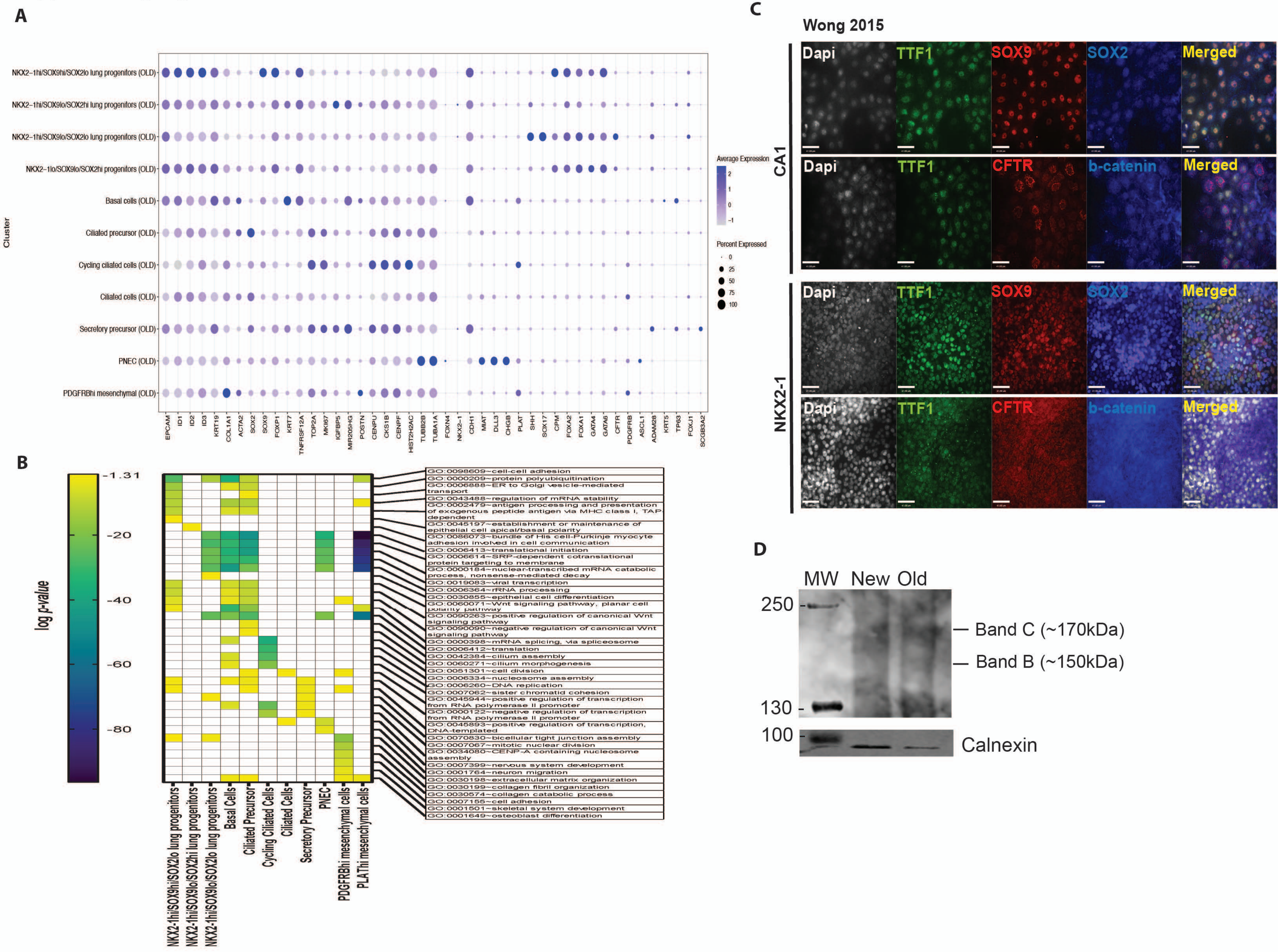

**Supplementary Fig. 4**

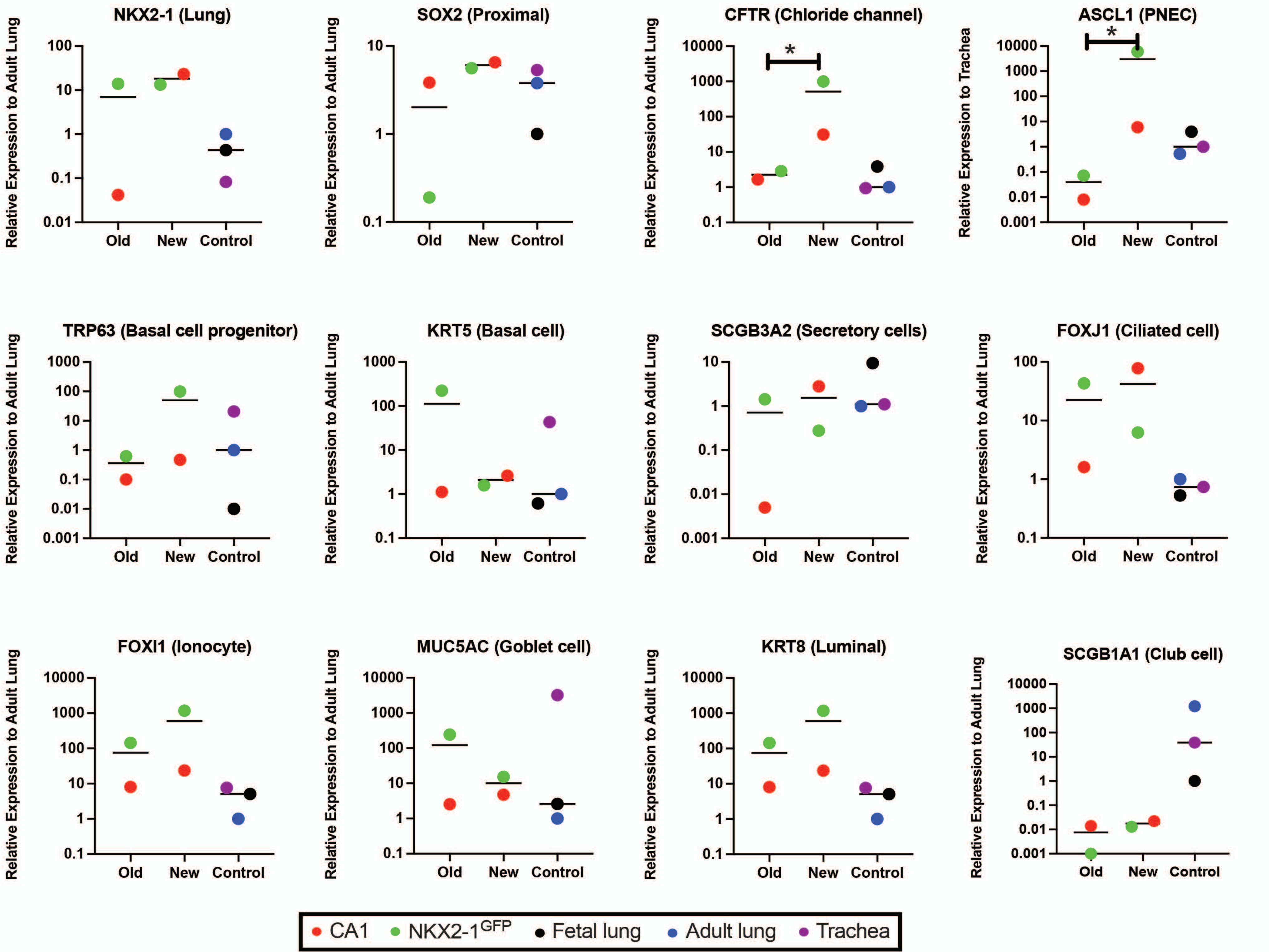

### Supplementary Fig. 5

A

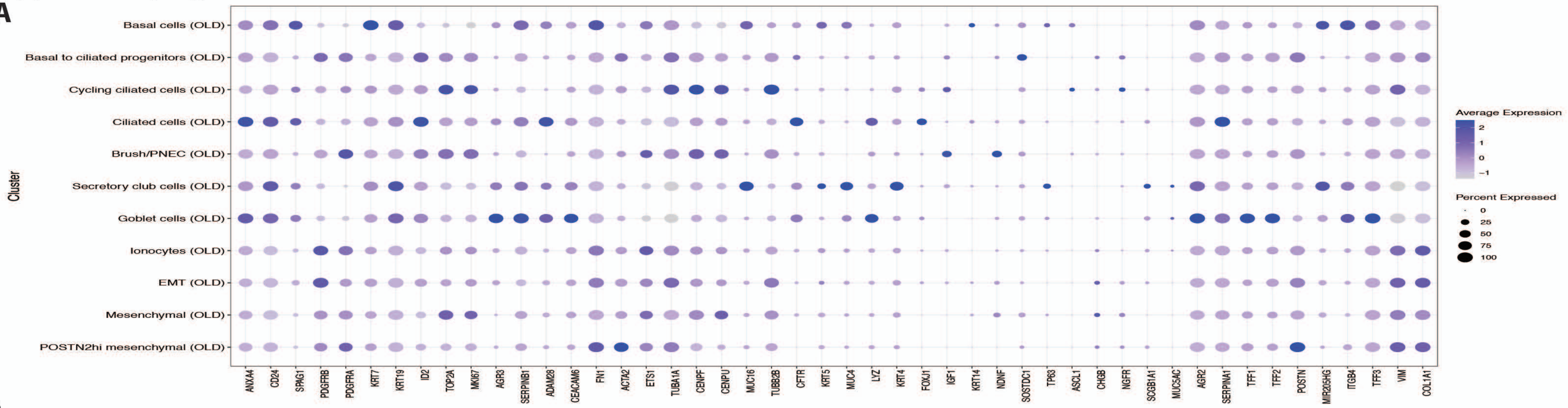


B

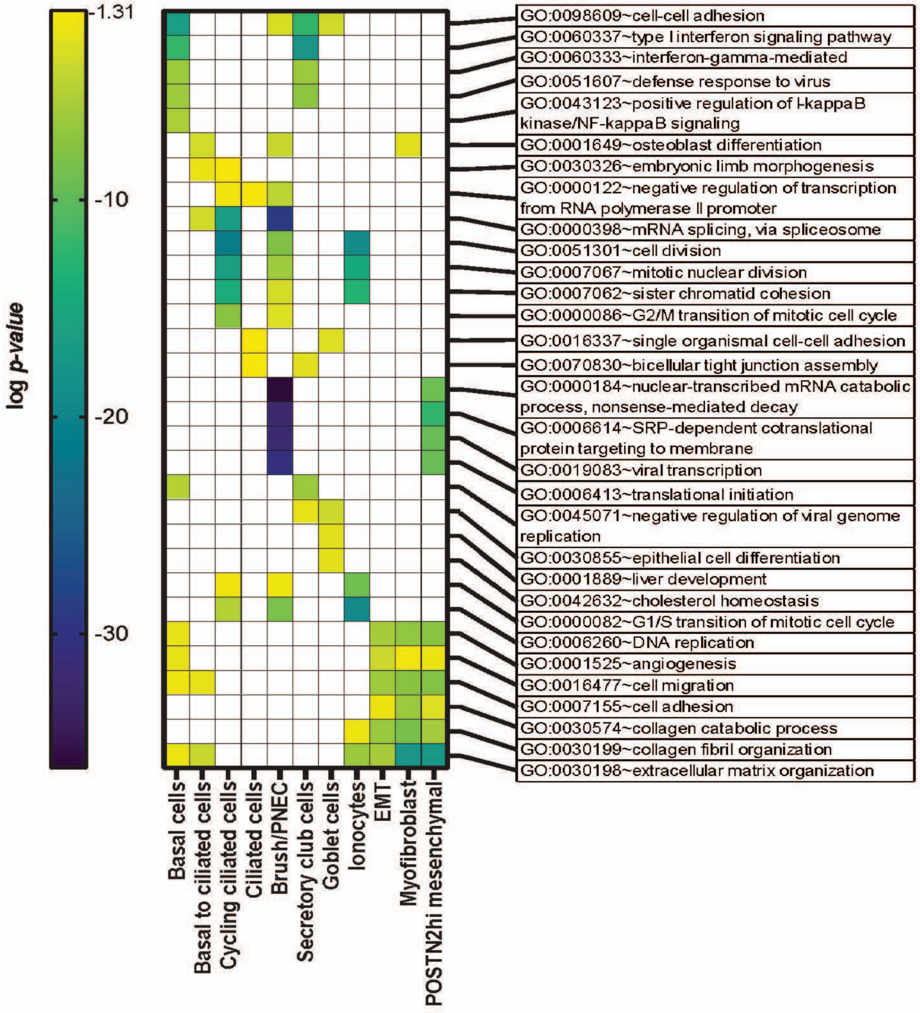
